# Fitness costs and benefits of gene expression plasticity in rice under drought

**DOI:** 10.1101/2021.03.16.435597

**Authors:** Simon C. Groen, Elena Hamann, Irina Ćalić, Colleen Cochran, Rachel Konshok, Michael D. Purugganan, Steven J. Franks

## Abstract

Genome-wide gene expression changes in response to environmental variability have been widely documented, but we lack detailed and comprehensive understanding of the interplay between this form of phenotypic plasticity and natural selection. Selection on expression plasticity may be limited by environment-specific costs, and plasticity may in turn affect selection on baseline expression levels. Here, we address this fundamental issue by measuring selection on drought-induced plasticity of leaf transcripts in field-grown rice populations. Selection disfavored switching off housekeeping genes under drought. This stress-induced dysregulation did not constrain selection on baseline transcript levels, suggesting compensatory evolution may be possible. Selection rarely acted strongly on individual transcripts but worked polygenically on gradual (continuous) plasticity of co-expressed gene modules regulating photosynthesis via known drought-responsive transcription factors. Finally, selection was tied to inefficient gene architectural features and metabolic costs of expression. Our study provides a genome-wide view of costs and benefits of gene expression plasticity.

## MAIN TEXT

### Introduction

Organisms often respond to spatial and temporal environmental variability through phenotypic plasticity. A trait is plastic when the same genotype expresses different phenotypic values across varying environments and canalized when this does not occur (Scheiner, 1993; Schlichting, 1986; Via and Lande, 1985). Examples of phenotypic plasticity include plants that produce inducible defensive compounds in response to herbivory (Agrawal et al., 2002; Groen et al., 2016a), and frogs that alter developmental timing in response to variation in food availability (Relyea, 2002a).

As rapid climatic changes continue, phenotypic plasticity may play an important role in species responses (Nicotra et al., 2010). Recent models suggest that plasticity could be advantageous if associated costs remain low (Scheiner et al., 2020), but plasticity might also hamper species’ evolutionary responses (Block et al. 2019; Ghalambor et al., 2015). The degree to which climatic changes cause plastic adjustments, and the relative importance of plasticity compared to other responses such as migration or evolution, are still debated (Merilä and Hendry 2014). While plasticity is ubiquitous, the extent to which plasticity is adaptive and evolving under selection is poorly understood (Arnold et al., 2019; Dudley and Schmitt, 1996; Hamann et al., 2016; Nussey et al., 2005; Pespeni et al., 2017; Saltz et al., 2018; Sultan, 2004; Van Buskirk and Steiner, 2009; Via et al., 1995), and whether plasticity is costly is also contentious (Auld et al., 2010).

Plasticity can be assessed for any quantitative phenotypic trait, but to date, most work has focused on higher-order traits such as growth or natural enemy defenses (Agrawal et al., 2002; Groen et al., 2016b; Relyea, 2002a,b; Schlichting and Levin, 1984; Valladares et al., 2006; Van Buskirk and Steiner, 2009). The expression level of a gene, however, is also a quantitative phenotypic trait, and variation in expression levels across environments constitutes plasticity, one with important links both to gene function and potentially to fitness (Groen et al. 2020). Because of advances in genomics technologies, it is now possible to obtain data on levels of gene expression throughout the genome for many individuals. It is thus possible to assess the degree of plasticity in gene expression (Figure 1A), selection on expression plasticity (Figure 1B), and whether plastic responses follow the same or opposite directions as selection on baseline expression—*i.e*., co-gradient or counter-gradient selection *sensu* Byars and colleagues (2007). Similarly, we can examine costs of expression plasticity (Figure 1B-C), as well as the nature of these costs, which provides important information on how individuals cope with environmental variation from the genetic to the phenotypic levels (Bittner et al., 2021; Kenkel and Matz, 2016; Mäkinen et al., 2016; Oostra et al., 2018). We currently lack a detailed, but at the same time comprehensive, assessment of the interplay between gene expression plasticity and natural selection, as previous studies either considered plasticity of the transcriptome as a unified whole (Bittner et al., 2021; Dayan et al., 2015; Ghalambor et al., 2015; Kenkel and Matz, 2016; Mäkinen et al., 2016; Oostra et al., 2018), only considered a small number if candidate genes (McCairns and Bernatchez, 2010), or measured selection on baseline expression levels, but not on expression plasticity (Ahmad et al., 2020; Koch and Guillaume, 2020).

**Fig. 1.**
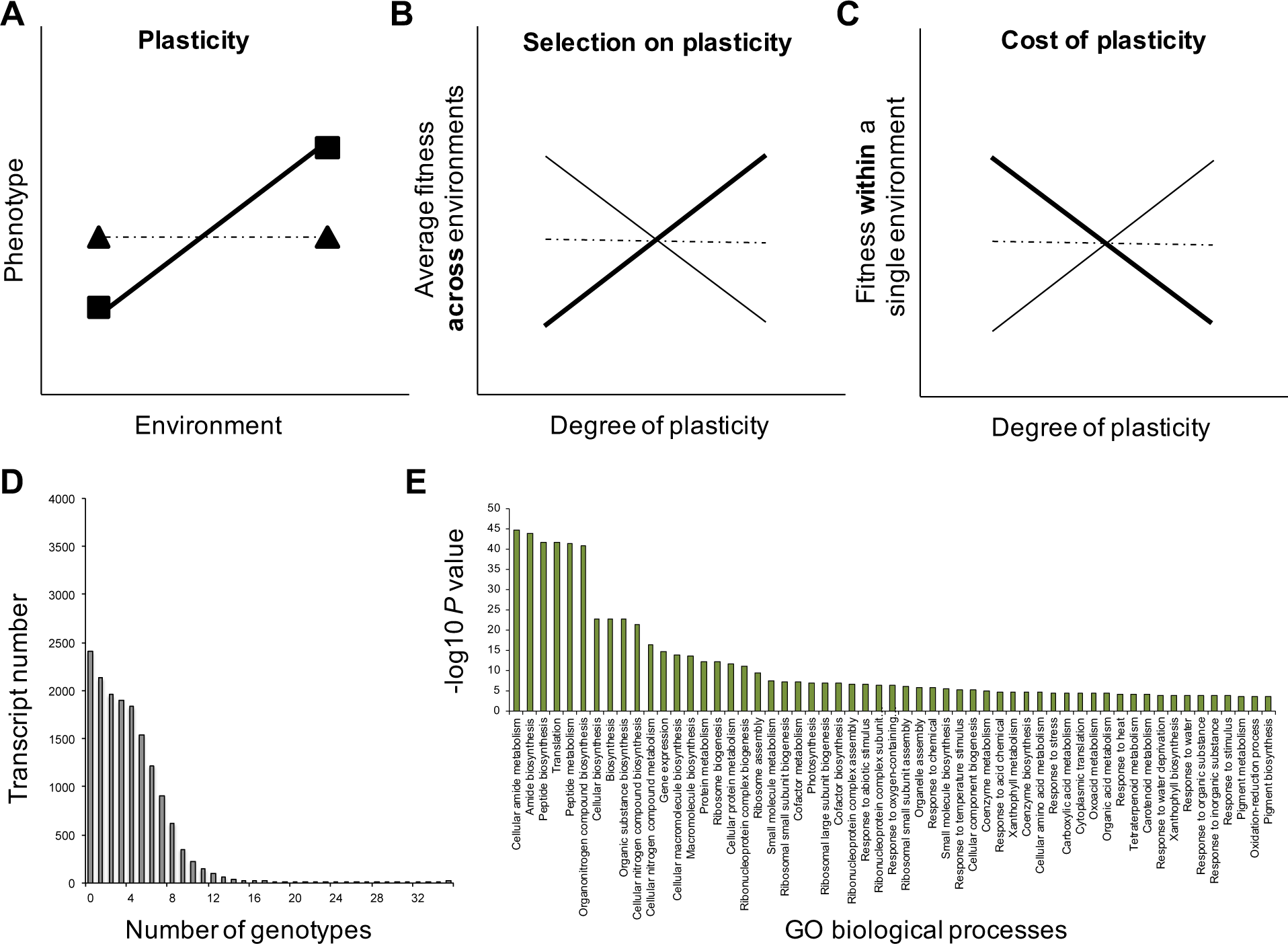
Analysis of selection on population-level heterogeneity in transcript level plasticity. **a,** The degree of phenotypic plasticity of genotypes as the slope of reaction norms with a plastic genotype (bold line) displaying different phenotypes across environments and a canalized (non-plastic) genotype (hatched line) maintaining the same phenotype across environments. **b,** Selection on plasticity can occur when environments are heterogeneous in space and time. When average fitness across environments increases with degree of plasticity (bold positive slope), plasticity is favored by natural selection and adaptive. In contrast, when average fitness across environments decreases with degree of plasticity (negative slope), selection would favor canalization (*i.e.*, loss of plasticity) and plasticity would be maladaptive. Plasticity can also be neutral and not under selection (hatched non-significant slope). **c,** Plasticity costs may constrain evolution of plasticity when genotypes with greater plasticity have lower fitness within a single environment (bold negative slope). However, when genotypes with greater plasticity have higher fitness (positive slope), or when the degree of plasticity does not affect fitness within environments (hatched non-significant slope), there is no evidence for plasticity costs that would constrain its evolution. **d,** Number of Indica genotypes in which each transcript is influenced significantly by drought (*P* value < 0.05) for each transcript identified from one-way ANOVAs. **e,** Gene ontology biological processes enriched in the tail of the distribution of differentially expressed transcripts, with each transcript influenced by drought in at least 10 genotypes (FDR *q* value<0.05).

In this study, we examined selection on plasticity of gene expression in rice, *Oryza sativa* (L.). Rice is an important crop and genetic model system, and can be sub-divided in two main varietal groups—Indica and Japonica—which we considered separately to account for population structure (Choi et al., 2020; Groen et al., 2020; Gutaker et al., 2020; McCouch et al., 2016; Wang et al., 2018). Rice landraces have been grown for millennia across gradients of factors such as soil moisture and temperature (Gutaker et al., 2020). Fluctuations within each of these environments occur and can be exacerbated by climate change, substantially influencing rice yields and fitness (Wing et al., 2018). We conducted a large-scale field experiment in which 216 Indica and Japonica accessions (mostly inbred landraces) were subjected to dry or wet conditions, and measured levels of gene expression and fitness. We previously described widespread variation in gene expression, as well as differences in selection on gene expression patterns between dry and wet environments (Groen et al., 2020).

In this study, we take advantage of this dataset to determine how selection and costs shape drought-induced gene expression plasticity and how plasticity in turn may affect selection on baseline expression levels in rice using quantitative, population and systems genetics. Specifically, we examine: 1) the heritabilities and sources of gene expression plasticity, 2) whether plasticity is under selection and/or associated with costs, 3) if plasticity constrains selection on baseline expression, 4) what factors impact patterns of selection on plasticity, and 5) which biological processes may be influenced by selection on expression plasticity.

### Results

#### Plasticity Is Heritable and Mostly Due to Rank-Order Changes in Reaction Norms

We previously observed significant plasticity (an effect of environmental drought) for ∼10% of rice transcripts, genetic variability in plasticity (G×E) for ∼90% of transcripts, and cross-environment heritability for nearly all transcripts (Groen et al., 2020). Here, we further characterized patterns of G×E interactions under drought (Figure 1A).

As observed for the model plant *Arabidopsis thaliana* (Des Marais et al., 2012), levels of many transcripts only responded to drought in one or a few accessions (Figure 1D; Supplementary Tables 1-2). The top ∼5% of most frequently differentially expressed transcripts among Indica accessions (771 transcripts with an environment effect in ≥10 accessions) was enriched for drought-related gene ontology (GO) biological processes, including “response to water deprivation” (Figure 1E; Supplementary Table 2). These patterns appeared to hold more generally, and drought-related processes were similarly enriched in the most frequently differentially expressed transcripts among Japonica accessions (Supplementary Figures 1A-B; Supplementary Tables 3-4). Expression plasticity heritabilities, at a median *H^2^* of ∼0.26, were lower than the heritabilities of baseline expression levels within each environment, at a median *H^2^* of ∼0.51, for Indica and Japonica (Figures 2A-B; Supplementary Figures 2A-B; Supplementary Tables 5-6). Furthermore, for both varietal groups, G×E interaction was mostly due to rank-order changes between reaction norms (Figures 2C-E; Supplementary Figures 2C-E; Supplementary Tables 5-6).

**Fig. 2.**
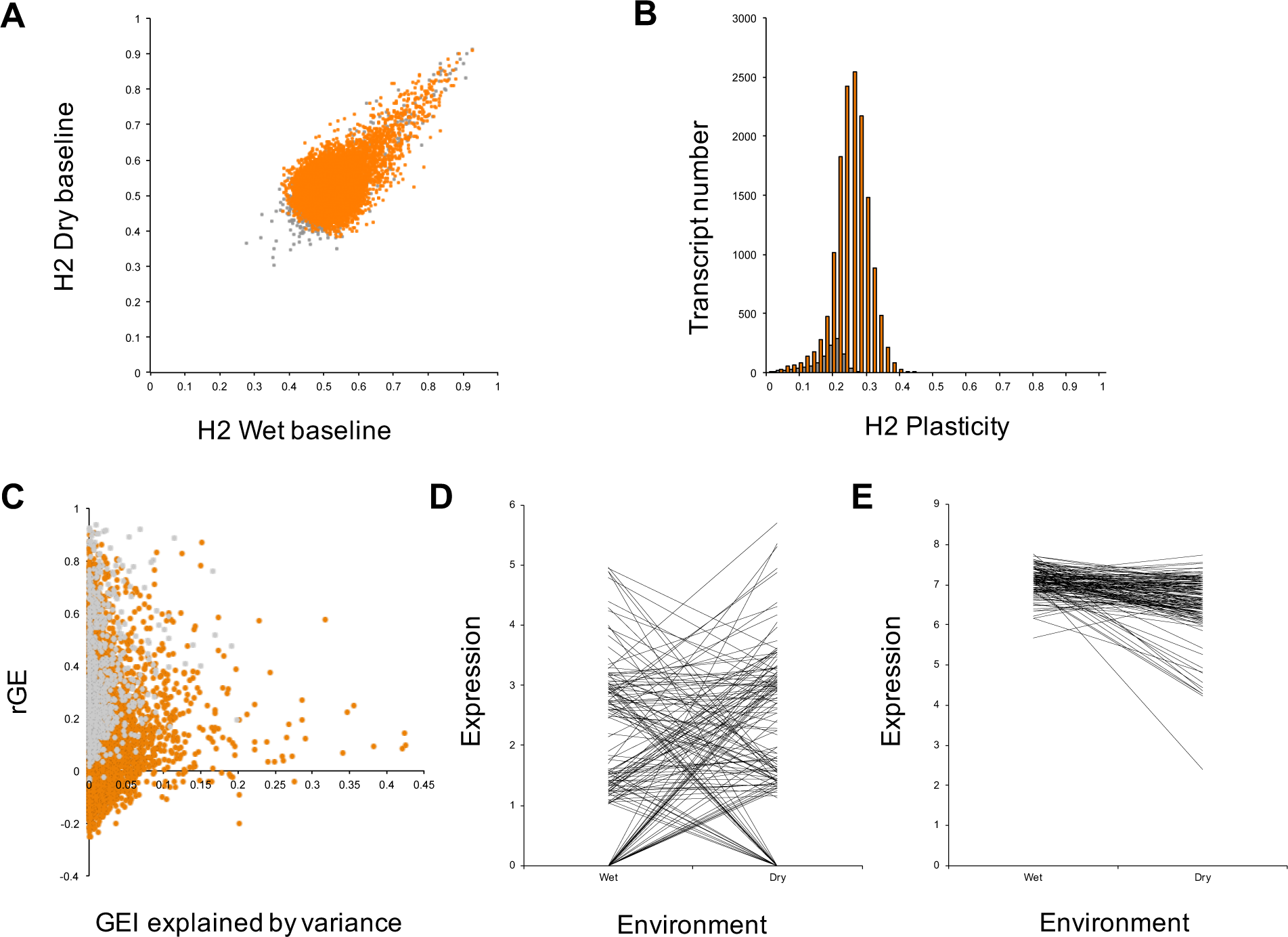
Heritability of transcript level plasticity. **a,** Bivariate plot of broad-sense heritability (*H^2^*) estimates for baseline transcript levels in the Indica populations in wet and dry conditions. Orange dots indicate significant G×E interaction variance (FDR *q* value<0.001), and grey non-significant. **b,** Distribution of *H^2^* estimates for transcript level plasticity across wet and dry conditions. Orange columns denote significant G×E interaction variance (FDR *q* value<0.001), and grey non-significant. **c,** Proportion of G×E interaction variance attributed to changes in among-accession variance in wet and dry conditions versus cross-environment genetic correlation. Orange dots indicate significant G×E interaction variance (FDR *q* value<0.001), and grey non-significant. **d,** Reaction norms of a transcript for which G×E interaction variance is mostly determined by deviation of cross-environment genetic correlation from unity, as indicated by abundant line crossing. **e,** Reaction norms of a transcript for which more G×E interaction variance is determined by changes in among-accession variance in wet and dry conditions, as indicated by less-abundant line crossing and wider among-accession variance in one environment than the other.

#### Known Drought-Responsive Transcripts Are Among the Most Plastic

Next, we divided transcripts in two categories according to prevailing patterns of expression level plasticity in the Indica and Japonica accessions. Transcripts either displayed graded (*i.e.*, continuous) variation in between-environment expression or discrete (*i.e*., on/off) states for some or all accessions, where expression was present in one environment and absent in the other (Stearns 1989).

Transcripts detected in at least two replicates for ≥75% of accessions in both environments, were classified as gradually plastic transcripts (GPTs) and as discretely plastic transcripts (DPTs) if detected above this threshold in one environment, but not the other. Applying these criteria, we analyzed 3,772 GPTs and 3,058 DPTs for Indica, as well as 3,789 GPTs and 3,580 DPTs for Japonica. Inclusion of transcripts within one or the other category was not influenced by general patterns of presence/absence variation (PAV) of genes in *O. sativa* (Figure 3; Supplementary Figure 3; Supplementary Tables 7-8).

**Fig. 3.**
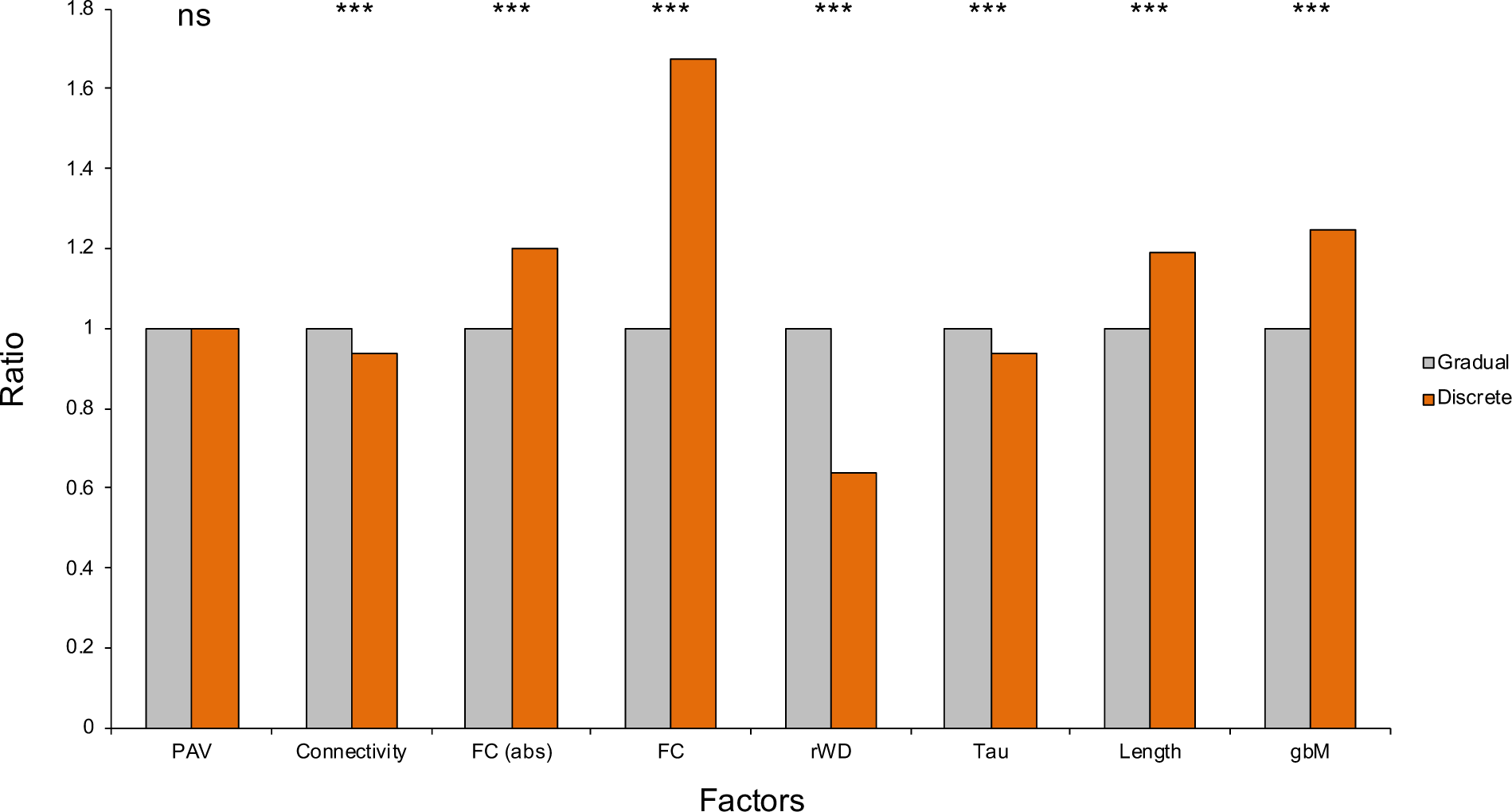
Structural and regulatory features of gradually and discretely plastic transcripts. Average levels of factors that characterize discretely plastic transcripts (DPTs) relative to average levels of these factors for gradually plastic transcripts (GPTs) in Indica. Values for the former have been normalized relative to the latter. PAV = presence/absence variation, FC = fold change, Tau = tissue specificity, gbM = gene body methylation, ns = non-significant; *** indicates *P*<0.001.

**Fig. 4.**
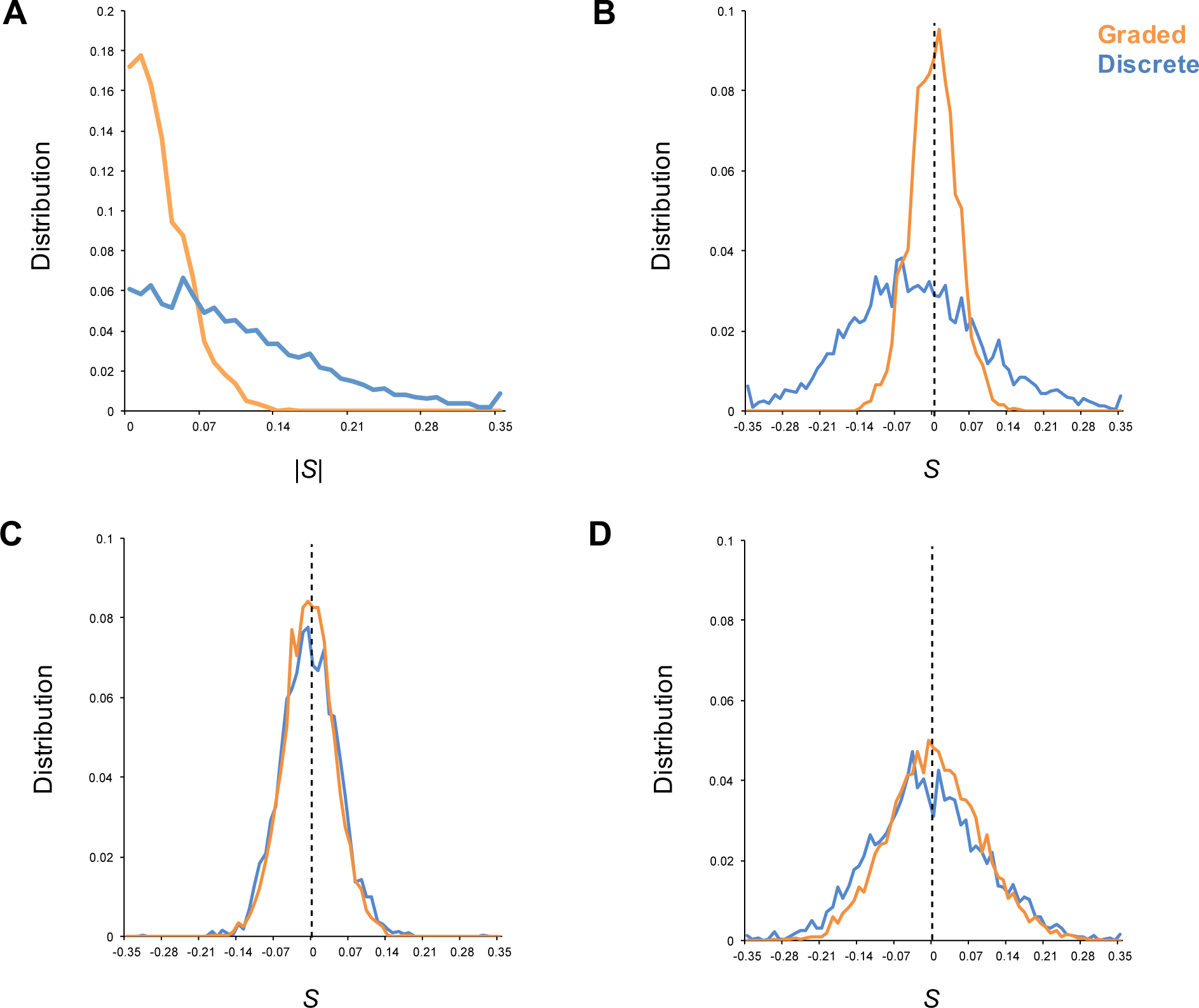
The strength and pattern of selection on expression plasticity differ between gradually and discretely plastic transcripts. **a,** The strength of selection |*S*| on gene expression plasticity in Indica when considering global across-environment (global) fecundity as fitness measure differed between gradually (orange) and discretely (blue) plastic transcripts (GPTs and DPTs, respectively). **b,** Positive directional selection on plasticity was stronger than negative directional selection for GPTs, and this pattern was inversed for DPTs. **c,** In wet conditions plasticity costs were generally stronger than benefits for GPTs. **d,** This pattern was inversed under drought. Selection did not show a significant bias towards costs for DPTs in either environment. See text for statistical information.

We then calculated plasticity levels for each Transcript×Genotype combination using a metric that was best suited to each category. We selected unsigned plasticity metrics to avoid biases regarding transcripts’ biological roles; for example, transcription factors can be activators or repressors (Wilkins et al., 2016). Levels of GPTs can be regarded as quantitative traits, and we calculated the simplified relative distance plasticity index (RDPI_s_) for normally distributed traits (Valladares et al., 2006). However, for non-normally distributed DPT expression (null for >25% of genotypes in one of the two environments), we calculated the coefficient of variation over the environments based on means (CV_m_), which is strongly correlated with RDPI_s_ (Supplementary Figure 4, Supplementary Tables 9-10; Valladares et al., 2006).

GPTs in the top 5% of RDPI_s_ values were enriched for drought-related GO biological processes, including “response to water deprivation” (Supplementary Figures 5A-B and 6A-B). In contrast, DPTs in the top 5% of CV_m_ values had no clear enrichment of such processes aside from ones related to protein transport and metabolism (Supplementary Tables 9-10). However, several known drought-related transcripts were present in the top 10 of CV_m_ values, including *OsRab16B*, coding for a dehydrin, and *OsPYL6*, involved in ABA perception, in Indica (Supplementary Figure 5C, Supplementary Table 9; Dey et al., 2016), and the transcriptional activator *ZFP182*, a regulator of drought responses, and *OsPP2C49*, also involved in ABA perception, in Japonica (Supplementary Figure 6C; Supplementary Table 10; Huang et al., 2012; Zong et al., 2016).

**Fig. 5.**
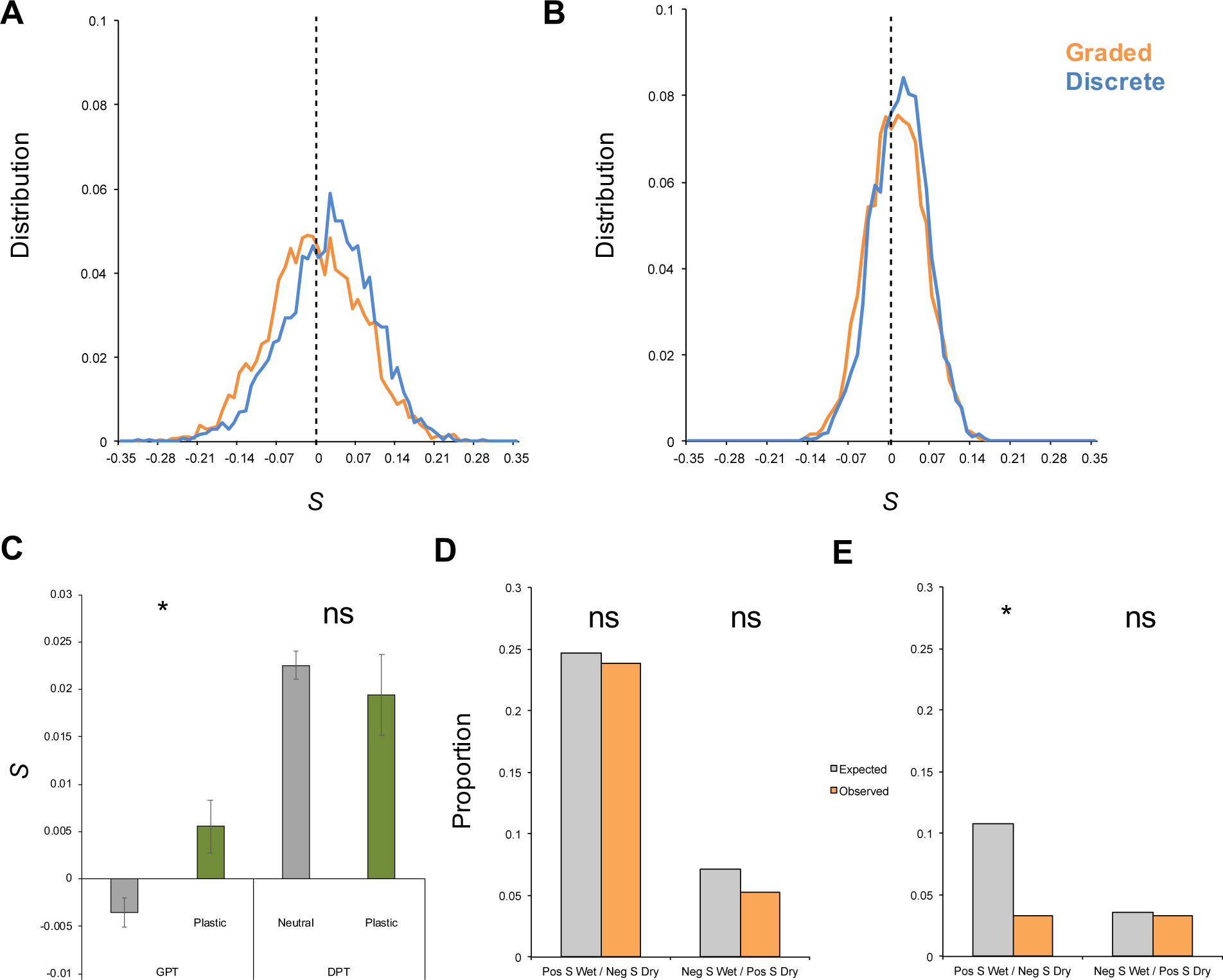
The strength and pattern of selection on baseline expression levels tend to be congruent between gradually and discretely plastic transcripts. **a,** Under drought selection tended to benefit higher baseline expression levels for both gradually and discretely plastic transcripts (GPTs and DPTs, respectively) in Indica. **b,** This pattern held true in wet conditions as well. See text for statistical information. **c,** Selection on baseline expression under drought was co-gradient with the levels of expression plasticity. **d,** Antagonistic pleiotropy is as common as expected by chance for GPTs. **e**, For DPTs, on the other hand, antagonistic pleiotropy is less common than expected among transcripts for which lower baseline expression levels are favored by selection under drought. Ns = non-significant; * indicates *P*<0.05.

#### Gradual Plasticity Is Frequently Favored by Selection

Having obtained an overview of the variation in genome-wide transcriptional plasticity among Indica and Japonica accessions, we then measured phenotypic selection on plasticity and plasticity costs (Figure 1B-C), taking into account baseline gene expression levels (Van Kleunen and Fischer, 2005). We calculated selection differentials that represent total (direct and indirect) selection by analyzing covariance between gene expression (measured as the genotypic average across replicate individuals of baseline transcript levels and between-environment plasticity in transcript levels) and fecundity fitness for each genotype as the average number of filled grains produced across replicates and environments (Lande and Arnold., 1983; Rausher, 1992). To assess potential costs of phenotypic plasticity, this selection analysis was repeated within environments. The idea is that a cost of plasticity is indicated when there is positive selection on plasticity when fitness is averaged across environments but negative selection on plasticity when considering fitness within an environment (Fig. 1B-C).

For GPTs in Indica, selection on expression plasticity appeared to be relatively weak. Transcriptome-wide selection on plasticity was |*S*|_median_=0.0291, with only ∼2.5% of transcripts showing |*S*|>0.1 (Figure 4A; Supplementary Table 11). This suggests that—for most genes— variation in expression plasticity is (nearly) neutral. Selection differentials showed a slight overall bias for stronger positive selection (greater fitness with greater plasticity) than for negative selection (lower fitness with greater plasticity) with *S*_median(pos)_=0.0302 and *S*_median(neg)_=−0.028, respectively (Mann–Whitney *U*-test [MWt], one-tailed *P*=0.0401; Figure 4B, Supplementary Table 11).

When plasticity costs occurred, they were generally higher in dry (|*S*|_median(dry)_=0.0581) than wet (|*S*|_median(wet)_=0.032) conditions (MWt, *P*<0.0001; Figures 4C-D, Supplementary Table 11). However, such costs occurred relatively more frequently in the wet environment (*S*_median(neg)_=|−0.033| > *S*_median(pos)_=0.030; MWt, *P*=0.03; Figure 4C, Supplementary Table 11) than in the dry environment (*S*_median(pos)_=0.0615 > *S*_median(neg)_=|−0.0547|; MWt, *P*=0.0004; Figure 4D, Supplementary Table 11).

Selection behaved similarly in Japonica (Supplementary Text, Supplementary Figure 7, Supplementary Table 12). Taken together, these results suggest that selection on plasticity of GPTs is generally more constrained by costs in the wet versus the dry environment.

#### Discrete Plasticity Is Generally Disfavored by Selection

Plasticity of DPTs was overall under stronger selection than that of GPTs in Indica with |*S*|_median(graded)_=0.0291 and |*S*|_median(discrete)_=0.0879, respectively (MWt, *z*=-42.16, *P*<0.0001; Figure 4A, Supplementary Table 13), indicating that the population distribution of plasticity levels for drought-responsive DPTs was further removed from the phenotypic optimum. DPTs showed lower connectivity than GPTs based on measurements over a time series of the tightness of their co-expression with other transcripts (0.93× difference, *P*=7.24×10^-36^; Figure 3; Plessis et al., 2015), and network buffering may be less effective in reducing effects of variation in their expression on fitness (Groen et al., 2020).

Contrary to selection on GPTs, *S* showed an overall bias for stronger negative than positive values for DPTs with *S*_median(pos)_=0.0746 and *S*_median(neg)_=−0.0973, respectively (MWt, *P*<0.0001; Figure 4B, Supplementary Table 13), indicating that plasticity of DPTs was generally selected against and canalization was beneficial. When plasticity costs occurred, these were generally higher in the dry than the wet environment for DPTs as for GPTs, with |*S*|_median(wet)_=0.036 and |*S*|_median(dry)_=0.069, respectively (MWt, *P*<0.0001; Figures 4C-D). However, again in contrast to patterns observed for GPTs, plasticity costs were not more prevalent in either the wet (*S*_median(pos)_=0.0363 ∼ *S*_median(neg)_=|−0.0357|; MWt, *P*=0.7114; Figure 4C, Supplementary Table 13), or the dry environment (*S*_median(pos)_=0.0683 ∼ *S*_median(neg)_=|−0.0693|; MWt, *P*=0.8572; Figure 4D, Supplementary Table 13).

In Japonica, selection on plasticity was generally also stronger for DPTs than GPTs (Supplementary Text, Supplementary Figure 7, Supplementary Table 14), indicating that the expression distributions for DPTs were overall further from adaptive peaks.

#### Discretely Plastic Transcripts Show Features of Expression Dysregulation

Next, we studied why selection impacted plasticity of DPTs and GPTs differently. In general, DPTs showed stronger environmental effects on expression than GPTs (1.2× difference in absolute log_2_ fold change, *P*=7.86×10^-18^; Figure 3, Supplementary Table 7), in particular stronger downregulation of expression in dry conditions (1.67× difference in log_2_ fold change, *P*=1.81×10^-18^; Figure 3), indicating DPT expression was more frequently shut down rather than activated under stress. Furthermore, when expressed, DPTs showed lower expression than GPTs, yet still higher than the expression of less regularly expressed transcripts (normalized log_2_ expression levels of 6>3.19>1.19 in wet conditions, and 5.86>2.96>1.07 in dry conditions for GPTs, DPTs and other transcripts, respectively; Supplementary Table 7). As expected because of their more discrete expression patterns, these transcripts exhibited weaker cross-environmental genetic correlations (0.64× difference, *P*=2.09×10^-61^; Figure 3).

One class of genes for which stress-induced repression might result in lower fitness is the class of housekeeping genes, which are essential for continued growth and development (Deprost et al., 2007). A first sign that many DPTs could have housekeeping roles is that they had wider potential expression across tissues as indicated by a 0.93× difference in the tissue specificity index τ (*P*=3.29×10^-24^; Figure 3; Yanai et al., 2005).

Housekeeping genes further tend to be longer, have fewer TF binding sites and TATA-boxes in their promoters, and more gene body methylation (gbM; Aceituno et al., 2008). Promoters of DPT-encoding genes did indeed contain fewer TF binding sites (median of 5 instead of the of 6 regulatory elements for GPTs, *P*<0.0001; Supplementary Figure 8A, Supplementary Table 15), and TATA-boxes (23.4% vs. 25.7%, *P*=0.002; Supplementary Figure 8A, Supplementary Table 15), and DPTs were longer than GPTs (1.19× difference, *P*=1.99×10^-41^; Figure 3, Supplementary Table 7), all in agreement with a housekeeping role. Furthermore, gbM levels were higher for the less abundant DPTs than for the more abundant GPTs (1.25× difference, *P*=8.12×10^-11^; Figure 3), matching previous observations on gbM and gene expression for field-grown rice (Wang et al., 2020). All of these patterns were similar overall for Japonica (Supplementary Figures 3 and 8B, Supplementary Tables 8 and 15).

Genes with high gbM levels show more consistent expression across cells (Horvath et al., 2019), which has contributed to the view that gbM has homeostatic functions for gene expression (Zilberman, 2017). Combined with the other pieces of evidence that an important part of DPTs may come from housekeeping genes (Figure 3, Supplementary Figure 3), the higher frequency of negative over positive *S* for the plasticity of DPTs, and their predominant pattern of downregulation under drought, our observations on DPTs could indicate stress-induced dysregulation of housekeeping genes and other important genes. This may indeed be maladaptive and have negative fitness consequences (Deprost et al., 2007; Kremling et al., 2018).

#### Selection on Baseline Expression Matched Selection on Expression Plasticity

Given the patterns of selection on transcript plasticity that we found, we had four predictions about patterns of selection on baseline expression within wet and within dry conditions. First, we expected to observe directional selection for higher baseline expression levels of DPTs under drought if their downregulation indeed constitutes dysregulation. This expectation was confirmed with *S*_median(pos)_=0.0624 being stronger than *S*_median(neg)_=−0.0472 (MWt, *P*<0.0001; Figure 5A, Supplementary Table 16). Second, given the result that plasticity of DPTs was generally selected against, we should observe a concomitant bias for higher baseline expression levels in wet conditions. This was again confirmed with *S*_median(pos)_=0.0411 and *S*_median(neg)_=−0.0267, respectively (MWt, *P*<0.0001; Figure 5B, Supplementary Table 16). Third, since selection tended to favor slightly higher baseline expression in wet conditions for a majority of transcripts in phenotypic selection analysis (Groen et al., 2020), we expected this pattern to be repeated for GPTs in genotypic selection analysis. This pattern indeed repeated itself with *S*_median(pos)_=0.0402 and *S*_median(neg)_=−0.0305, respectively (MWt, *P*<0.0001; Figure 5B, Supplementary Table 16). Fourth, given our observations that selection showed a mild bias towards higher plasticity for GPTs, and tended to favor a slightly higher baseline expression under drought for a majority of transcripts in a previous phenotypic selection analysis (Groen et al., 2020), we also expected selection to promote higher baseline GPT expression under drought. A pattern of slightly stronger positive than negative selection with *S*_median(pos)_=0.0577 and *S*_median(neg)_=−0.0558, respectively, was in keeping with this expectation (MWt, *P*<0.0001; Figure 5A, Supplementary Table 16).

Signatures of selection on baseline expression levels in Japonica generally matched those in Indica, except for a general pattern of negative rather than positive selection on baseline expression levels of GPTs under drought, with *S*_median(pos)_=0.0805 and *S*_median(neg)_=−0.0952, respectively (MWt, *P*<0.0001; Supplementary Figure 9, Supplementary Table 17).

#### Plasticity and Pleiotropy Minimally Constrain Selection on Gene Expression

Having found effects of selection on gene expression plasticity, we then examined whether plasticity may constrain selection on baseline expression levels under drought through testing if a relation was co- or counter-gradient (Byars et al., 2007; Conover and Schultz, 1995; Eckhart et al., 2004; Koch and Guillaume, 2020). We also tested if selection on baseline expression under drought may be constrained by antagonistic pleiotropy with respect to selection on expression in wet conditions (Anderson et al., 2013).

Among DPTs, we observed no significant patterns of co- or counter-gradient selection between plasticity and *S* for baseline expression levels in dry conditions. However, among GPTs, we found that plasticity was significantly co-gradient with selection for higher baseline expression levels under drought (Figure 5C, Supplementary Table 18). This indicated that plasticity does not appear to have strong effects on limiting selection on baseline expression under drought in Indica and may even invigorate selection.

Patterns of antagonistic pleiotropy between environments were no different from that expected by chance for GPTs in Indica (Figure 5D, Supplementary Table 18). However, in keeping with a deleterious effect of stress-induced expression downregulation for many DPTs, we observed significantly fewer instances of antagonistic pleiotropy than expected by chance that were linked to negative *S* on baseline expression levels under drought (Figure 5E). This suggested that for DPTs selection for higher baseline expression levels under drought tended to counteract deleterious downregulation of gene expression under drought.

In Japonica, we observed no biases in patterns of antagonistic pleiotropy, although we did observe a pattern of counter-gradient selection on plasticity that matched the general pattern of negative rather than positive selection on baseline expression levels of GPTs under drought (Supplementary Figure 9, Supplementary Table 19).

#### Selection on Drought-Induced Discrete Plasticity Is Shaped by Metabolic Costs

Drought not only limits water availability, but also access to soluble nutrients such as phosphorus and nitrogen. It further constrains carbon fixation, since photosynthesis requires water. Transcription of mRNAs and their translation into protein rely on these nutrients and production may be strained when supply is limited (Acquisti et al., 2009). Baseline expression and expression plasticity might thus incur metabolic costs. And this is not the only source of costs; plasticity costs may further be genetic in nature such as when genetic pleiotropy occurs (DeWitt et al., 1998).

We compared how transcripts with positive and negative *S* in the five separate selection scenarios we studied (plasticity across environments plus plasticity costs and baseline expression within each environment) differed for 25 characteristics, including metabolic features such as N content and ATP requirements of gene expression and protein production, gene architectural features such as transcript and protein length and the number and size of introns, as well as network features such as transcript connectivity as a proxy for pleiotropy.

Starting with selection on plasticity of DPTs in Indica, transcripts showing positive *S* had longer primary transcripts (1.06× difference, *P*=0.002), with higher total ATP and N requirements (Figure 6A, Supplementary Table 20), than transcripts for which selection promoted canalization (showing negative *S*). Transcripts with positive *S* further had longer 5’ UTRs and showed higher expression and plasticity levels (*≥*1.03× difference, *P≤*0.009; Figure 6A), again indicative of patterns of co-gradient selection (Byars et al., 2007). The proteins these transcripts encoded were longer (1.12× difference, *P*=0.001), with higher total requirements in terms of C, N and ATP (Figure 6A). Continuing with plasticity costs of DPTs in wet conditions, we observed that the importance of most of these factors disappeared, except for CDS length (*≤*0.99× difference, *P*=0.016; Figure 6A). However, under drought, plasticity costs were characterized by primary transcript length (1.06× difference, *P*=0.004; Figure 6A), 5’ UTR GC content (0.98× difference), and transcript connectivity, our proxy for pleiotropy (1.03× difference, *P≤*0.004; Figure 6A).

**Fig. 6.**
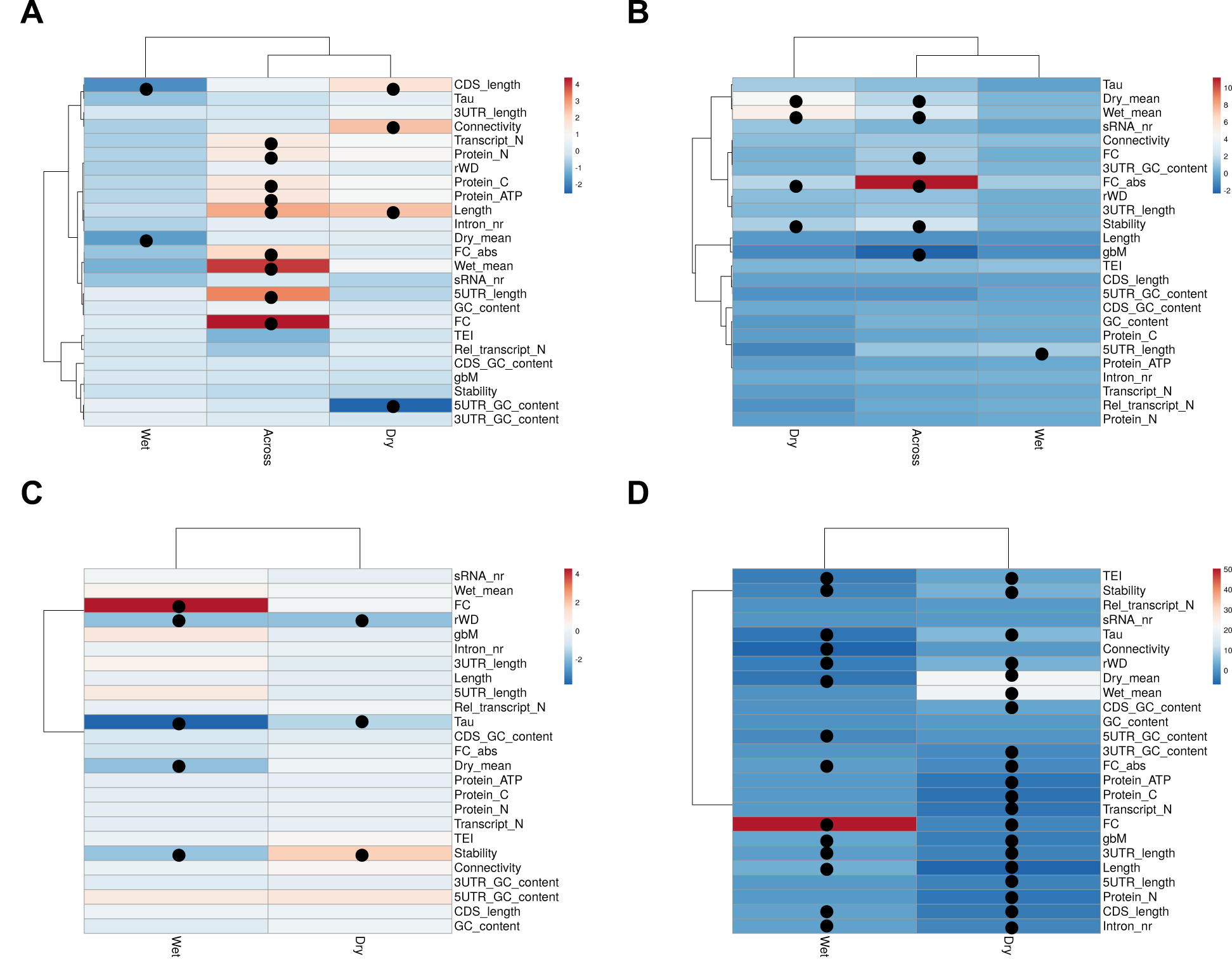
Comparison of defining characteristics of transcripts with positive and negative selection differentials on expression plasticity and baseline expression in the Indica populations. Gradually and discretely plastic transcripts (GPTs and DPTs, respectively) were compared for 25 architectural, biochemical and network characteristics in the five selection scenarios we studied for DPTs and GPTs separately: across- and within-environment selection on plasticity for DPTs **(a)** and GPTs **(b)**, as well as selection on baseline expression in wet and dry conditions for DPTs **(c)** and GPTs **(d)**. Panels depict the negative log10 of the *P* values of the *t*-tests between the transcripts with positive and negative *S*. Positive changes indicate a characteristic was more strongly associated with transcripts defined by positive *S* than with transcripts defined by negative *S*, while negative changes indicate the opposite pattern.

#### Selection on Drought-Induced Gradual Plasticity Is Shaped by Gene Architecture

Analysis of selection on plasticity of GPTs revealed transcripts with positive *S* had higher expression and plasticity (*≥*1.02× difference, *P≤*0.021; Figure 6B, Supplementary Table 20) than transcripts for which selection promoted canalization (showing negative *S*), just as we observed for DPTs (Figure 6A). These transcripts were also less stable (1.02× difference, *P*=0.007; Figure 6B), showing that transcript secondary structure may be important (Su et al., 2018). Finally, transcripts with positive *S* came from loci with lower gbM (0.87× difference, *P*=0.005; Figure 6B), in keeping with a previously observed link between gbM and adaptive gene expression plasticity in corals (Dixon et al., 2018). For plasticity costs of GPTs, none of these factors were highly significant in wet conditions (Figure 6B). However, except for gbM and transcript stability, which were no longer highly significant, transcript level and plasticity did shape plasticity costs under drought as they had done for patterns of selection (*≥* 1.04× difference, *P≤*0.02; Figure 6B).

The importance of gene architecture, expression level and plasticity, as well as metabolic costs, were generally visible for selection on plasticity in Japonica as well (Supplementary Text, Supplementary Figure 10, Supplementary Table 21).

#### Selection on Baseline Expression Under Drought Further Minimizes Costs

Given our observation that selection tended to favor plasticity of more highly expressed and plastic transcripts, and among DPTs also favored plasticity of transcripts that require more resources to produce and translate, we next investigated if similar factors were important in shaping selection on baseline expression levels.

In wet conditions, selection favored discrete plasticity of transcripts that were more strongly down-regulated (1.44× difference), and had lower cross-environment genetic correlation and expression under drought (*≤*0.98× difference, *P≤*0.016; Figure 6C, Supplementary Table 20). DPTs with positive *S* showed less tissue-specific expression and higher transcript stability (*≤*0.98× difference, *P≤*0.017; Figure 6C). In dry conditions most of these patterns were similar, although not highly significant (Figure 6C).

For GPTs, more factors appeared to be important than for DPTs. In wet conditions, GPTs under positive selection for higher baseline expression showed lower connectivity and less tissue-specific expression (*≤*0.97× difference, *P≤*4.002×10^-5^; Figure 6D, Supplementary Table 20), in keeping with previous selection analyses on gene expression (Groen et al., 2020). Furthermore, these GPTs were longer as primary transcripts, with longer CDSs, contained more introns, and were more plastic (*≥*1.06× difference, *P≤*0.011; Figure 6D). In particular, they were more strongly downregulated under stress (13.81× difference), and had lower cross-environment genetic correlations (0.89× difference, *P≤*0.0006; Figure 6D). GPTs with positive *S* further have lower translation efficiency (*≤*0.99× difference), while showing higher stability (0.98× difference), and are transcribed from loci that have stronger gbM (1.15× difference, *P≤*0.009; Figure 6D). Transcript-specific negative and positive correlations between mRNA stability and translation in plants have been observed previously when they experienced either wet or dry conditions (Kawaguchi et al., 2004).

Under drought, the influence of several of these factors reversed in direction. GPTs under positive selection for higher expression in dry conditions were more tissue-specific and had higher translatability (*≥*1.04× difference), while being less stable (1.03× difference), containing fewer introns (0.98× difference), and not just being shorter as primary transcripts, but also containing shorter 5’ UTRs, 3’ UTRs and CDSs (*≤*0.92× difference, *P≤*0.005; Figure 6D). Given these observations, and that shortness has been linked to higher translatability previously (Zhao et al., 2017), it is not surprising that transcripts with positive *S* had lower N requirements as well (0.91× difference, *P≤*5.09×10^-5^; Figure 6D). Furthermore, these transcripts were less strongly downregulated under drought than transcripts with negative *S* (0.78× difference; *P≤*0.012; Figure 6D). GPTs with positive *S* also showed higher cross-environment environment genetic correlations (1.14× difference) and were transcribed from genes with lower gbM (0.85× difference, *P≤*0.0008; Figure 6D).

The proteins encoded by GPTs under positive selection for higher baseline expression levels under drought were not only shorter, but also had lower total requirements in terms of C, N and ATP (*≤*0.93× difference, *P≤*4.19×10^-5^; Figure 6D). Patterns were mostly similar for Japonica (but see Supplementary Text, Supplementary Figure 10, Supplementary Table 21). Taken together, these observations suggest that, overall, selection may favor lower baseline expression, but higher plasticity levels, of metabolically costly transcripts and proteins, as well as more efficient gene architectures. Furthermore, the importance of genetic costs of plasticity such as through pleiotropy appears to be more limited, since transcript connectivity (a proxy of the potential for pleiotropy) was linked with benefits rather than costs of discrete plasticity under drought (Figure 6A).

#### Discrete Switch-Off of Translation-Related Transcripts Under Stress Is Selected Against

Prior studies showed that some of the most energy-costly housekeeping processes of the cell such as mRNA translation, which in plants is to some extent regulated by the hormone ABA, are downregulated in times of stress (Baena-González, 2010; Deprost et al., 2007). Accordingly, we observed transcripts related to ABA perception and ABA responses among the most plastic DPTs (Supplementary Figure 5C, Supplementary Figure 6C). These processes contribute to a strong reduction or a halt in growth and development and could lead to an early senescence (Baena-González, 2010; Deprost et al., 2007). Indeed, in stressful conditions, translation is one of the first targets of regulation, and following even a mild drought a global decrease in translation can be measured (Kawaguchi et al., 2004).

To infer the biological relevance of selection on gene expression plasticity, we intersected the DPTs and GPTs with our analyses of plasticity heritability, since only significantly heritable heterogeneity will be changed in offspring. For Indica and Japonica, 2,139 and 2,075 DPTs had significantly heritable plasticity, respectively (Supplementary Tables 22-23). Using these DPTs, we aimed to test for correlated expression patterns across genes, summarizing expression across co-expression modules and improving power to detect selection (Hämälä et al., 2019; Huang et al., 2020). Specifically, we examined whether some modules had stronger average selection on gene expression plasticity than others. Yet, connectivity of DPTs proved to be too weak for the detection of co-expression modules (Figure 3, Supplementary Figure 3). Nonetheless, we obtained a picture of the processes potentially targeted by selection by analyzing enriched GO biological processes among transcripts with negative and positive *S*. We only considered processes enriched for both Indica and Japonica to enhance the likelihood that signatures of selection are biologically meaningful. Both varietal groups shared enrichment of processes related to translation, growth, and flowering among DPTs with negative *S* on expression plasticity, meaning selection favored decreased plasticity in genes related to these functions. Among transcripts with positive *S* (selection for increased plasticity), we observed enrichment for processes related to ABA responses (Figure 7A, Supplementary Figure 11A, Supplementary Table 24). No individual transcript made a stringent Bonferroni threshold, but three came within an order of magnitude (*P*<0.0001; Supplementary Text, Supplementary Tables 22-23), including *OS09T0505800-01* in Indica, for which plasticity was selected against (*S*=-0.42, *P*=5.68×10^-5^; Supplementary Text, Supplementary Table 22). This transcript codes for a uridine kinase involved in pyrimidine metabolism and may play key roles in photosynthesis and abiotic stress responses (Dong et al., 2019b).

**Fig. 7.**
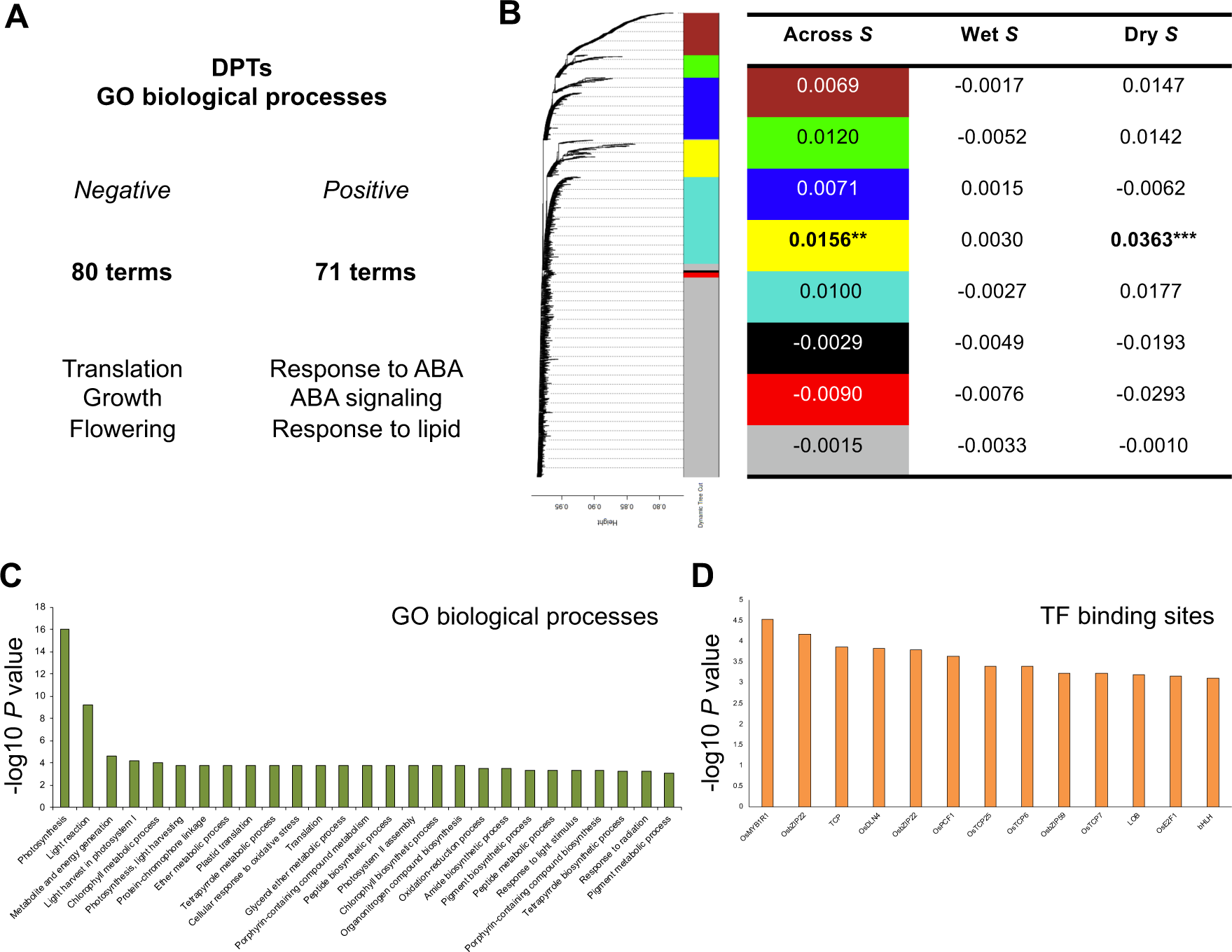
Selection may have polygenic effects on plasticity levels of transcript co-expression modules. **a,** Gradually plastic transcripts (GPTs) in Indica are enriched for different Gene Ontology (GO) biological processes among transcripts with positive and negative selection differentials on their plasticity levels. **b,** Selection acts more strongly on one module of co-expressed GPTs than on other modules. See text for statistical information. **c,** The module under selection is enriched for photosynthesis-related biological processes. **d,** In addition, the gene promoters of the module’s transcripts show enrichment of binding sites for bZIP, MYB, and TCP transcription factors.

Plasticity costs appeared non-significant for individual DPTs, since none of the selection differentials came within an order of magnitude from the Bonferroni threshold (*P*>0.0002; Supplementary Tables 13-14). Combined with the analyses of the distributions of selection differentials, this suggests that the maladaptive nature of plasticity in DPT expression is more apparent at a polygenic level.

#### Selection Favors Graded Plasticity of Photosynthesis- and Stress-Related Transcripts

2,240 and 1,740 GPTs had significantly heritable plasticity in Indica and Japonica, respectively (Supplementary Tables 25-26), and grouped into co-expressed transcript modules using WGCNA. For Indica, 1,238 transcripts yielded seven modules (Figure 7B; Supplementary Table 25). Yellow module 4 showed elevated levels of positive selection on expression plasticity compared to other modules and transcripts with unique expression patterns (*S*=0.0156, *P*=0.003). This module was enriched in photosynthesis-related GO biological processes (Figure 7C; Supplementary Table 25), suggesting polygenic selection on expression plasticity of photosynthesis-related genes. There was no evidence of plasticity costs for any of the modules in wet conditions (yellow module 4: *S*=0.003, *P*=0.162; Figure 7B), and under drought *S* for expression plasticity of yellow module 4 was even more strongly positive than across environments (*S*=0.0363, *P*<0.0001; Figure 7B). Thus, adjusting photosynthesis to fluctuating water availabilities appeared to have particularly beneficial fitness consequences.

Gene promoters for the transcripts in yellow module 4 were enriched for binding sites of several transcription factors central to an environmental gene response network derived from plants grown in the same field site (Plessis et al., 2015; Wilkins et al., 2016). These transcription factors include OsbZIP22, OsE2F1, and OsMYB1R1, which are involved with regulating expression levels of photosynthesis-related genes in response to heat stress and dehydration (Figure 7D; Wilkins et al., 2016). This overlap emphasizes the regulatory coherence of yellow co-expression module 4 and the biological relevance of selection on expression plasticity of this module.

One individual transcript in particular showed significant selection on expression plasticity, even following Bonferroni correction (Supplementary Table 25). This transcript was *OS09T0556400-01* from *OsPHT4;5*, coding for an inorganic phosphate transmembrane transporter. *OsPHT4;5* is differentially expressed in response to abiotic stress (Ruili et al., 2020), and plastic expression of phosphate transporters contributes to optimization of shoot growth across stressful and non-stressful conditions (Dong et al., 2019a).

Six other transcripts showed selection on expression plasticity that came within an order of magnitude from the Bonferroni cut-off (Supplementary Table 25). Three encoded metabolite transporters for either glucose, sphingolipids, or malate, and experienced positive selection (*S≥*0.132, *P≤*0.0011). Vacuolar malate uptake and storage via tonoplast dicarboxylate transporters allows plants to make osmotic adjustments and regulate stomatal opening and closing (El Mahi et al., 2019; Emmerlich et al., 2003). Two other transcripts displayed negative *S* (*i.e*., selection for canalization; *S≤*-0.125, *P≤*0.0008). One of these, from *OsISC5*/*OsISU2*, coded for an iron-sulfur cluster assembly protein, which is not only involved in iron homeostasis and the maintenance of essential housekeeping processes such as respiration and DNA repair, but also in photosynthesis and abiotic stress responses (Kesawat et al., 2012; Liang et al., 2014).

In Japonica, there were two modules that showed stronger selection on expression plasticity than other modules and transcripts with unique expression patterns (Supplementary Text, Supplementary Figure 11, Supplementary Table 26). Similar to our findings for Indica, among these modules we observed enrichment for photosynthesis- and stress response-related GO biological processes, and for promoter binding sites of transcription factors OsbZIP22, OsE2F1, and OsMYB1R1. Overall, these results suggest selection acted on expression plasticity of GPTs related to photosynthesis and stress responses, which is fitting in the context of drought response.

### Discussion

Our study has deepened our understanding of gene expression plasticity, in particular how it is affected by and in turn shapes natural selection. Taking a genome-wide overview of our findings for the plasticity of gene expression as thousands of molecular quantitative traits expression plasticity appears to share strong similarities with the more extensively studied plasticity of functional traits that are likewise quantitative in nature.

First, work in maize and *Drosophila melanogaster* showed that trait plasticity typically has lower heritability than baseline trait levels (Kusmec et al., 2017; Scheiner and Lyman, 1989). Similarly, in our study, the heritability of expression plasticity was generally lower than the heritability of baseline expression. Second, work on functional traits in Drosophila, Arabidopsis, and *Caenorhabditis elegans* established that the mechanistic underpinnings of G×E interaction are mostly due to rank-order changes in reaction norms (Carreira et al., 2013; Gutteling et al., 2007; Ungerer et al., 2003), and we found this to be the case for a majority of transcripts as well.

Additionally, the similarities between trait and gene expression plasticity extended further to the manner in which selection acted on levels of plasticity. Selection on expression plasticity was (nearly) neutral in most transcripts examined, a pattern that resembles the distribution of selection differentials for functional trait plasticity across hundreds of traits in a wide variety of plant and animal populations (Hendry, 2016; Van Buskirk and Steiner, 2009). Yet, how often gene expression plasticity is adaptive or non-adaptive is still debated. Several previous studies considered genome-wide gene expression plasticity and arrived at opposite conclusions about the adaptiveness of plasticity when the transcriptome was considered as a unified whole (Bittner et al., 2021; Dayan et al., 2015; Ghalambor et al., 2015; Kenkel and Matz, 2016; Mäkinen et al., 2016; Oostra et al., 2018). While these groundbreaking studies did much to deepen our understanding of the adaptive nature of gene expression plasticity, our fine-grained approach enabled us to pinpoint individual transcripts and classes or modules of transcripts for which selection either favored or disfavored plasticity.

Indeed, our genotypic selection analyses allowed us to observe that plasticity was not only favored by selection for a majority of GPTs in Indica, but that it was also co-gradient with selection for higher baseline transcript levels under drought, and that these transcripts were enriched for regulating known drought-responsive biological processes. This finding was in keeping with our previous observation that Indica accessions with broader drought-induced changes in gene expression experience greater fitness in dry conditions (Groen et al., 2020). Transcripts with beneficial plasticity were further enriched for growth- and photosynthesis-related processes, which were shown to have fitness benefits in several functional genetic studies (*e.g*., Kromdijk et al., 2016; Yoon et al., 2020). Interestingly, family members of some of the transcription factors involved, such as MYB1R, were previously also found to play an important role in rapid evolutionary response to contemporary drought episodes in *Brassica rapa* (Hamann et al. 2020).

In contrast, the majority of DPTs appeared to experience selection for more stringent canalization and mostly showed selection against expression plasticity. This pattern was likely the result of stress-induced dysregulation of gene expression (Kremling et al., 2018), with a subset of housekeeping genes, such as ones involved in mRNA translation, being downregulated to the point of non-expression in the dry environment. Importantly, discrete plasticity was generally not counter-gradient to selection on baseline expression levels, which suggests that expression of these transcripts is not constrained by plasticity and that changes compensatory to non-adaptive plasticity of these transcripts have a higher chance of evolving.

Lastly, we were able to identify likely sources of plasticity costs, which have been notoriously difficult to detect (Auld et al., 2010; Van Kleunen and Fischer, 2005, 2007). While we did not observe a prominent role for pleiotropy in exerting plasticity costs (Figures 5D-E and 6A), we did find roles for metabolic costs and costs of inefficient gene architectural features in shaping selection on expression plasticity as well as baseline expression. Selection appeared to act towards minimizing metabolic costs, in particular favoring discrete drought-induced plasticity of metabolically costly transcripts and proteins (Figure 6B), and lower baseline expression of GPTs with high metabolic requirements under drought (Figure 6D). These observations are in keeping with those from earlier studies on costs of gene expression in evolution experiments with *Escherichia coli* bacteria (Frumkin et al., 2017). Selection further favored gradual expression plasticity of genes with higher mRNA turnover (Figure 6B), linking fitness with mRNA structural features (Su et al., 2018).

Patterns of selection on baseline expression levels matched the manner with which selection acted on expression plasticity and further acted towards minimizing metabolic costs of gene expression under stress. The fact that we observed parallel patterns of selection in two independent sets of rice populations from the Indica and Japonica varietal groups bolsters the general relevance of our observations.

### Materials and Methods

We used representative studies from the literature for determination of sample size (Des Marais et al., 2012; Gutteling et al., 2007; Huang et al., 2020; Lafuente et al., 2018; Ungerer et al., 2003; West et al., 2007). Individuals of each genetic identity were allocated randomly to the wet and dry environments and their planting order was randomized as laid out in an alpha lattice design. Investigators were blinded to the genetic identity of individuals when sampling, processing samples and assessing results.

#### Data sources

We previously assessed transcriptome variation in two rice *Oryza sativa* populations—one consisting of 136 varietal group ‘Indica’ accessions (comprising the indica and *circum*-aus subgroups) and the other of 84 varietal group ‘Japonica’ accessions (comprising the japonica and *circum*-basmati subgroups)—in a field experiment in the Philippines (Groen et al., 2020).

Identical sister populations, with three individuals per accession as biological replicates, were planted in a continuously wet paddy and a field that imposed intermittent drought (Groen et al., 2020). We used 3′-end mRNA sequencing to measure transcript levels in leaf blades of the 1,320 plants at 50 days after sowing, corresponding to 17 days after withholding water in the dry field. Genetic variation was observed in the levels of 15,635 widely expressed transcripts and partitioned variance among genetic, environmental and interactive (G×E) effects.

For each gene-expression trait (*i.e*. transcript), following an ANOVA approach, a mixed-effect general linear model was fit, which included a term for “accession” or “genotype” (G) as a random factor, “field environment” (E) as a fixed factor, the G×E interaction as a random factor, and the error variance (ε). Cross-environment genetic correlations were estimated as *r_GE_* = *cov_ij_* / *σ_i_σ_j_*, where *cov_ij_* is the covariance of accession means between a trait as *i* in the wet and *j* in the dry field environments, and *σ_i_* and *σ_j_* are the square roots of the among-genotype variance components for the trait in the wet and dry field environments. For the analyses described below we consider fecundity, quantified as the numbers of filled grains produced (Groen et al., 2020), as our fitness proxy.

#### Differential gene expression analyses

To obtain estimates of population heterogeneity in gene expression plasticity, we performed targeted ANOVAs for each accession individually by fitting a fixed-effect general linear model, including a term for “field environment” (E) as a fixed factor, and the error variance (*ε*). The significance of the variance explained by the environment factor was tested using an *F*-test. Population-level variation in expression plasticity was quantified as the number of accessions with significant drought-modified expression for each of the transcripts at FDR *q*<0.05 (Des Marais et al., 2012).

#### Heritability and genotype×environment interaction analyses

We estimated broad-sense heritabilities per environment for transcripts by estimating the amount of variance explained by genotype within each environment as *H^2^* = *σ^2^_G_* / (*σ^2^_G_* + *σ^2^_E_*) where *σ^2^_G_* is the among-genotype variance component and *σ^2^_E_* is the error variance (West et al., 2007). For the heritability of plasticity, we used *H^2^* = *σ^2^_GE_* / *σ^2^_P_*, where *σ^2^_GE_* is the genotype-environment interaction (G×E) variance component and *σ^2^_P_* is the total phenotypic variance (Lafuente et al., 2018; Scheiner and Lyman, 1989).

Significant G×E interaction may come from two sources: deviation of the cross-environment genetic correlation *r_GE_* from unity, and differences in the among-genotype variance between environments. We determined the contribution from each of these sources using the following equation (Gutteling et al., 2007; Ungerer et al., 2003): *V_G×E_* = 0.5 (*σ_i_* -*σ_j_*)^2^ + *σ_i_ σ_j_* (1 - *r_GE_*), where *V_G×E_* is the G×E interaction variance component, *σ_i_* and *σ_j_* are the square roots of the among-genotype variance components for the trait in the wet and dry field environments, and *r_GE_* is the cross-environment genetic correlation.

#### Plasticity analyses

Based on how widespread transcripts were expressed across genotypes, individual replicates, and wet and dry environments, we selected two sets of transcripts: 1) gradually plastic transcripts (GPTs – continuous expression) were expressed by at least two individual replicates of >75% of genotypes in a population in each of the two environments, 2) discretely plastic transcripts (DPTs –expression switches on/off) had to show the same widespread expression, but in only one environment.

GPTs can be considered similar to quantitative functional traits, and we quantified the plasticity of transcripts levels across environments for each genotype as the simplified relative distance plasticity index (RDPI_s_), which allows for statistical comparisons of genotypes (Valladares et al., 2006). We calculated RDPI_s_ as the absolute difference of mean genotypic trait values across environments divided by the mean genotypic trait value in the wet environment, following Valladares et al. (2006).

For DPTs, whose expression is null in one of the two environments, we calculated the coefficient of variation over the environments based on means (CV_m_) as in Schlichting and Levin (1984) and Schlichting (1986), a measure strongly correlated with RDPI_s_ (Valladares et al., 2006).

#### Genotypic selection analyses

To examine the strength and pattern of selection on gene expression and expression plasticity, we conducted genotypic selection analyses based on the genotypic value averaged across replicate individuals (Rausher, 1992). Fitness for each genotype was the average number of filled grains produced across replicates. Genotypic selection analyses were conducted in three contexts: genotypic values across the wet and dry environments, as well as within the wet and dry environments.

We retained genotypes for analysis only when all three replicates in each environment had a positive number of filled grains. This resulted in 87 genotypes for the Indica and 31 for the Japonica population, with each genotype representing a data point in the regressions of standardized transcript expression or transcript plasticity on relative fitness. Relative fitness was calculated for each genotype by dividing the average number of filled grains produced across replicates by the mean number of filled grains produced by all genotypes. For the analysis of GPTs, a few individuals did not express a particular transcript, and we excluded genotypes from analysis when fewer than two replicates expressed a transcript in each environment. For the analysis of DPTs, we excluded genotypes from analysis when fewer than two replicates expressed a transcript in both environments. Mean trait values were then standardized to a mean of 0 and a standard deviation (s.d.) of 1 across and within environments, respectively.

For GPTs plasticity is estimated through RDPI_s_ defined as: *P_j_* = | *Z_j,k=2_* –*Z_j,k=1_* | / *Z_j,k=1_* ; for DPTs plasticity is estimated through CV_m_ defined as: *P_j_* = s.d. (*Z_j,k=2_*, *Z_j,k=1_*) / mean (*Z_j,k=2_*, *Z_j,k=1_*), where *j* is Genotype, *k* is the Focal environment, *Z* is the Transcript value, and *P* is the Transcript plasticity. The plasticity index values for each trait or transcript were standardized across environments to a mean of 0 and s.d. of 1.

First, we estimated standardized linear selection differentials *S* as the regression coefficient of the standardized mean trait value on relative fitness (Conner and Hartl, 2004; Lande and Arnold, 1983; Relyea, 2002b). Second, to examine selection on transcript plasticity, we estimated *S* as the partial regression coefficient of the standardized mean transcript level value and of the transcript plasticity index on relative fitness (Conner and Hartl, 2004; Lande and Arnold, 1983; Relyea, 2002b; Van Tienderen, 1991), using the following formula: *W_j_*_,*k*_ = Constant*_k_* + *α_k_ * Z_j_*_,*k*_ + *β_k_ * P_j_*, where *W* is Mean fitness, *j* is Genotype, *k* is the Focal environment, *Z* is the Transcript value, *P* is the Transcript plasticity, *α* is the selection differential on Transcript value, and *β* is the selection differential on Transcript plasticity.

To assess potential costs of phenotypic plasticity, this genotypic selection analysis was repeated within environments. The idea is that a cost of plasticity is indicated when there is positive selection on plasticity when fitness is averaged across environments but selection against plasticity when fitness is determined within an environment. For example, a highly plastic genotype might experience relatively greater fitness across environments than a non-plastic genotype, but this highly plastic genotype might show low fitness relative to the non-plastic genotype within the drought environment, which would indicate a cost of this plasticity. Here, we used relative fitness calculated as the mean genotypic filled grain number produced in one environment (wet or dry) divided by the average filled grain number of all genotypes in that same environment. In addition, we performed separate genotypic selection analyses to obtain selection differentials on baseline transcript expression within each environment using the formula: *W_j_*_,*k*_ = Constant*_k_* + *α_k_ * Z_j_*_,*k*_.

All genotypic selection analyses were performed in *R* v. 3.6.3 (R Core Team, 2016). Regression outputs were retrieved using the *broom* package in the *tidyverse* modeling set (Wickham et al., 2019), and we extracted selection differentials (*S*) for baseline transcript expression (*α)*, transcript plasticity (*β)*, and their corresponding statistical significance levels. Given the high number of transcripts analyzed, Bonferroni correction on *P* values to account for multiple testing would be highly conservative, and the analyses we were conducting were considered exploratory. Thus, we consider two-tailed *P* values for the gene expression genotypic selection analyses significant when a *P*-value comes within an order of magnitude of the Bonferroni cut-off at *α* = 0.05 / *n* in each respective analysis, unless otherwise indicated.

To assess co-gradient or counter-gradient selection (Byars et al., 2007), we first filtered those GPTs and DPTs that fell within the ∼10% of transcripts with a significant effect of environment on their expression as determined previously (Groen et al., 2020). Then, we analyzed whether significant differential expression in the dry versus the wet environment was linked to stronger positive or negative selection on baseline transcript levels under drought with two-sample Welch’s *t*-tests, controlling for unequal sample sizes and variances, in *R* v. 3.6.3 (*R* Core Team, 2016).

#### Factors affecting selection on gene expression plasticity

For a more in-depth view of the differences between GPTs and DPTs and to identify factors linked to the strength and pattern of phenotypic selection on gene expression and plasticity therein, we performed analyses considering factors previously found to influence evolutionary change in gene expression including cross-environment genetic correlation and heritability, primary transcript length, GC content, tissue-specificity, expression polymorphism and noise, grand mean expression level, transcript connectivity, as well as the number of *cis*-regulatory elements and in-degree for each transcript’s gene (Groen et al., 2020).

These were supplemented with data on gene body methylation, the number of transcripts, transcription start sites (TSSs) and introns for each gene, the presence or absence of a TATA-box in the promoter, presence/absence variation (PAV) in whole genes between different accessions in populations, transcript stability and translatability, and the length and GC content of 5’ UTR, 3’ UTR and CDS (Aceituno et al., 2008; Castillo-Davis et al., 2002; Frumkin et al., 2017; Zhao et al., 2017). Data on gene body methylation in non-stressed conditions were obtained from Chodavarapu *et al*. (2012), as processed by Elhaik *et al*. (2014), on transcripts and number of introns per gene from the Ensembl Plants BioMart release 43 *O. sativa* Japonica IRGSP-1.0 dataset (https://plants.ensembl.org/biomart/martview), on length and GC content of 5’ UTR, 3’ UTR and CDS, as well as translation efficiency, from an analysis of ribosome-associated mRNAs (Zhao et al., 2017), on transcript stability from an analysis of the *O. sativa* “structurome” (Su et al., 2018), on the number of TSSs and the presence/absence of a TATA-box from TSSPlant (Shahmuradov et al., 2017), on gene PAV among Indica and Japonica varietal group accessions from the 3K-RG project (Wang et al., 2018), and on codon usage from the Kazusa database (Nakamura et al., 2000).

Based on the codon usage data we calculated the total and relative (to molecule length) number of N atoms used per transcript (Kelly, 2018), and the number of C and N atoms used per protein (Arnold and Nikoloski, 2014). We also calculated the ATP expenditure per transcript using general eukaryote-derived values (Lynch and Marinov, 2015; Wagner, 2005), and the ATP expenditure for amino acids per protein using plant-derived values from Arabidopsis, ignoring the negligible protein assembly costs (Arnold and Nikoloski, 2014).

Data for each factor were grouped separately for DPTs and GPTs, and for transcripts with negative and positive selection differentials within these. We then conducted separate two-sample Welch’s *t*-tests in *R* v. 3.6.3 (*R* Core Team, 2016) for DPTs and GPTs, respectively, controlling for unequal sample sizes and variances, and using transcripts with negative and positive selection differentials as samples within each of the following five scenarios: selection on expression plasticity, costs of expression plasticity in wet conditions and in dry conditions, and selection on baseline expression in wet conditions and in dry conditions. Results were visualized in heatmap format with *ClustVis* (Metsalu and Vilo, 2015).

#### Co-expression analysis

To infer the biological relevance of selection on gene expression plasticity we selected GPTs and DPTs with significant plasticity heritability. We applied the following filters that ensured we considered relevant transcripts: 1) the G×E term had to be significant (FDR *q* value <0.001 for Indica, and FDR *q* value <0.01 for Japonica, respectively; Groen et al., 2020); 2) the cross-environment genetic correlation had to be <0.3, since transcripts with non-significant G×E had a median *r_GE_* of 0.3 - accordingly, heritably plastic transcripts should generally have a lower *r_GE_* (Groen et al., 2020); 3) differential expression had to reach a significance threshold of *P*<0.05 in at least four genotypes in the Indica population (multiplicatively breaching the Bonferroni threshold of *P*<0.05 / 3,772 transcripts =1.33×10^-5^, since 0.05^4^ ∼6.25×10^-6^), and three in the smaller Japonica population. The latter threshold was based on the most numerous class of transcripts in the Indica population, which are GPTs. A concern here would be that if transcripts are differentially expressed in all genotypes, then there would be no, or only a small, G×E effect. However, no transcript was differentially expressed in more than 78 genotypes out of the 136 in the Indica population, and 36 out of the 84 in the Japonica population, respectively, which is consistent with a G×E effect for all transcripts that passed our filtering steps. This left 2,240 and 1,740 GPTs as well as 2,139 and 2,075 DPTs with significantly heritable plasticity in the Indica and Japonica populations, respectively.

We constructed transcript co-expression networks through weighted gene co-expression network analysis (WGCNA) using the *WGCNA R* package available at https://horvath.genetics.ucla.edu/html/CoexpressionNetwork/Rpackages/WGCNA/ (Langfelder and Horvath 2008). We inferred separate networks for GPTs and DPTs with significantly heritable plasticity in the Indica and Japonica populations using expression data that was averaged per genotype within the wet and dry environment. For each of the four separate network construction analyses, data were used to define a co-expression similarity matrix using the absolute Pearson correlation coefficient between pairwise transcript profiles. The co-expression similarity based on GPTs was transformed into connection strengths (i.e., creating an adjacency matrix) by raising values to a soft thresholding power of 2 for Indica and 1 for Japonica, respectively, which best approximate scale-free topologies (model fit >0.8). For DPTs, there was no soft thresholding power that ensured approximation of scale-free topology, and we did not construct co-expression networks based on DPTs. For the GPTs, we determined clusters of transcripts with highly correlated expression across samples (*i.e.* modules) based on topological overlap dissimilarity measure in conjunction with linkage hierarchical clustering using the *blockwiseModules* function. Co-expressed transcripts were merged into modules of *≥*10 transcripts.

#### Gene-set enrichment analysis

For transcripts with evidence of selection on their baseline expression or plasticity levels (*i.e.* expression and plasticity patterns that significantly co-varied with plant fitness), we examined gene functional annotations. We hypothesized that if selection may act on gene expression plasticity, then the functional categories of these genes would include biological processes related to stress response and flowering time, which are associated with drought responses (Groen et al., 2020; Hamann et al., 2020).

We performed genes set enrichment analyses using the ShinyGO v0.61 gene-set enrichment tool (http://bioinformatics.sdstate.edu/go/) at default parameter settings (Ge et al., 2020). A Fisher’s exact test was applied to identify the most enriched KEGG terms, and Biological Process, Molecular Function, and Cellular Component GO terms.

#### Annotation of transcripts under selection

For individual transcripts under selection for their levels of plasticity, we performed further in-depth biological characterization. To assess functions of their genes, we examined GO terms for biological processes assigned from the UniProt database, and GO terms were grouped under functional categories using DAVID 6.8 (Huang et al., 2009a,b).

## Data Availability

The RNA-seq data are available from the Sequence Read Archive (SRA) under SRA BioProject accession number PRJNA588478. Processed RNA expression count data can also be found in Zenodo (https://zenodo.org/record/3533431 with DOI 10.5281/zenodo.3533431), alongside a sample metadata file with a key to the RNA sequence data in SRA BioProject accession number PRJNA588478. Data on gene expression and fecundity alongside the quantitative genetics analyses have been made available as Supplementary Material of an article we published previously (Groen et al., 2020).

## Acknowledgements

We thank New York University High Performance Computing for supplying computational resources. We are grateful to current and former members of the Franks and Purugganan laboratories for insightful discussions. This work was funded in part by grants from the Zegar Family Foundation and the NYU Abu Dhabi Research Institute to M.D.P., the National Science Foundation (DEB-1142784) to S.J.F., and (IOS-1546218 in the Plant Genome Research Program) to S.J.F. and M.D.P., a fellowship from the Gordon and Betty Moore Foundation/Life Sciences Research Foundation through Grant GBMF2550.06 to S.C.G., a fellowship from the Swiss National Science Foundation (P2BSP3_168833) to E.H., and a FCRH Summer Research Grant and Fordham-NYU Research Internship to C.C.

## Author contributions

S.J.F. and M.D.P. conceived and directed the project; E.H., S.C.G., I.Ć., S.J.F. and M.D.P. designed selection analyses; S.C.G., E.H. and I.Ć. processed fitness, functional trait and gene-expression data; E.H., C.C. and R.K. performed selection analyses; S.C.G. and E.H. performed statistical analyses; S.C.G., E.H., S.J.F. and M.D.P. wrote the manuscript, and all authors edited the manuscript.

## Supplementary Information

Supplementary figures and tables are available below.

## Supplementary Text

### The Majority of Gradually Plastic Transcripts Do Not Show Plasticity Costs in Japonica

Selection on gradual plasticity was generally stronger in Japonica than in Indica with |*S*|_median(Jap)_=0.1075 vs. |*S*|_median(Ind)_=0.0291 (Mann–Whitney *U*-test [MWt], *P*<0.0001; Supplementary Figure 7A, Supplementary Table 12), and while the overall bias for slightly stronger positive selection was similar to the bias in Indica with *S*_median(pos)_=0.1342 and *S*_median(neg)_=−0.1241, respectively, this bias was not significant (MWt, *P*=0.484; Supplementary Figure 7B, Supplementary Table 12).

In contrast to the Indica populations, we found that for Japonica costs and benefits of expression plasticity were generally highly magnified in the wet over the dry environment with |*S*|_median(wet)_=0.1265 and |*S*|_median(dry)_=0.0794, respectively (MWt, *P*<0.0001; Supplementary Figures 7C-D, Supplementary Table 12). In the wet environment, we did not detect plasticity costs for the majority of GPTs with *S*_median(pos)_=0.1381 and *S*_median(neg)_=−0.1188, respectively (MWt, *P*<0.0001; Supplementary Figure 7C, Supplementary Table 12). In the dry environment, this pattern was similar, and also here, positive selection differentials on expression plasticity were stronger overall than negative selection differentials: *S*_median(pos)_=0.0829 and *S*_median(neg)_=−0.077, respectively (MWt, *P*=0.0053; Supplementary Figure 7D, Supplementary Table 12). The latter is in full agreement with our findings for Indica.

### The Majority of Discretely Plastic Transcripts Do Show Plasticity Costs in Japonica

Selection on discrete plasticity, as with gradual plasticity, was generally also stronger in Japonica than in Indica with |*S*|_median(Jap)_=0.3035 vs. |*S*|_median(Ind)_=0.0879 (MWt, *P* < 0.0001; Supplementary Figure 7A; Supplementary Table 14). Furthermore, as with GPTs, in Japonica selection did not show a significant bias for positive or negative selection on discrete plasticity either, with *S*_median(pos)_=0.311 and S_median(neg)_=−0.3 (MWt, *P*=0.5029; Supplementary Figure 7B, Supplementary Table 14).

Like we observed for GPTs, in Japonica the costs and benefits of discrete plasticity were generally also larger in the wet environment than in the dry environment with |*S*|_median(wet)_=0.1462 and |*S*|_median(dry)_=0.1014, respectively (MWt, *P*<0.0001; Supplementary Figures 7C-D). And again, in the wet environment, we did not observe plasticity costs for the majority of DPTs with *S*_median(pos)_=0.1592 and *S*_median(neg)_=−0.138, respectively (MWht, *P*<0.0001; Supplementary Figure 7C, Supplementary Table 14). This time, the pattern was reversed in the dry environment, and negative selection differentials on discrete plasticity were stronger overall than positive selection differentials: *S*_median(pos)_=0.0848 and *S*_median(neg)_=−0.1099, respectively, indicative of plasticity costs (MWt, *P*<0.0001; Supplementary Figure 7D, Supplementary Table 14).

### Selection on Drought-Induced Discrete Plasticity Is Shaped by Gene Architecture in Japonica

In Japonica we found that, compared to DPTs with negative *S* for plasticity, DPTs with positive *S* had shorter primary transcripts and CDSs (*≤*0.94× difference), and these had lower total requirement for N (0.95× difference, *P≤*0.044; Supplementary Figure 10A, Supplementary Table 21). DPTs with positive *S* further were more GC rich, and had higher translation efficiency (*≥*1.01× difference, *P≤*0.008; Supplementary Figure 10A). In wet conditions, costs were associated with a bias for lower translatability among DPTs with negative *S*, and these transcripts were now also targeted by small RNAs at a lower density (*≥*1.08× difference, *P≤*0.037; Supplementary Figure 10A). However, in dry conditions the total N requirements per transcript still mattered, and DPTs with negative *S* incurred higher costs (0.99× difference, *P ≤* 0.047; Supplementary Figure 10A). In addition, higher drought-induced plasticity was now important (1.32× difference), which could be influenced by higher transcript stability (0.97× difference, *P≤*0.0043; Supplementary Figure 10A).

### Selection on Drought-Induced Gradual Plasticity Is Also Shaped by Metabolic Costs in Japonica

For GPTs our analyses showed that, in comparison with transcripts with negative *S* for plasticity, transcripts with positive *S* had lower expression (0.97× difference), and stronger directional drought-induced expression plasticity (1.41× difference, *P≤*0.014; Supplementary Figure 10B, Supplementary Table 21). For baseline expression and expression plasticity these patterns were opposite to what we observed for GPTs in Indica (Figure 6B). Transcripts with positive *S* were also targeted less often by small RNAs (0.71× difference), and showed lower GC content in the 3’ and 5’ UTRs (*≤*0.99× difference, *P≤*0.043; Supplementary Figure 10B), which may contribute to these patterns of regulation. Finally, transcripts under positive selection for plasticity had more tissue-specific expression (1.02× difference, *P*=0.04; Supplementary Figure 10B). This shows that these transcripts experience fewer constraints than transcripts under selection for canalization.

Plasticity costs for GPTs in wet conditions were linked with higher transcript levels (0.98× difference, *P*=0.042; Supplementary Figure 10B). Under drought on the other hand, the effects of weaker directional drought-induced expression plasticity on transcripts with negative *S* came to the fore again (1.35× difference, *P*=0.003; Supplementary Figure 10B). Interestingly, transcripts showing evidence of plasticity costs had lower N requirements than transcripts with positive *S* (1.06× difference), and the same pattern was visible with respect to requirements for N, C, and ATP that the proteins these transcripts encode incur (*≥*1.05× difference, *P≤*0.026; Supplementary Figure 10B).

Overall, it appears that selection may affect many of the same architectural, metabolic and regulatory features of transcripts in Japonica as in Indica, although the patterns of selection may sometimes differ because the populations start out at different distributions relative to the phenotypic optimum.

### Selection Strongly Alters Costs of Baseline Expression Under Drought in Japonica

Next, we checked how selection may influence transcript features related to baseline expression levels in Japonica. In wet conditions, selection favored higher baseline expression of DPTs with lower cross-environment genetic correlations (0.86× difference, *P*=0.012; Figure 7C, Supplementary Figure 10C, Supplementary Table 21). Other characteristics showed opposite influence in dry vs. wet conditions: for example, transcripts downregulated under drought experienced selection for higher baseline expression in dry conditions (0.74× difference), but for lower baseline expression in wet conditions (0.94× difference, *P≤*0.002; Supplementary Figure 10C).

For GPTs more factors appeared to be important than for DPTs. In wet conditions, GPTs under positive selection for higher baseline expression showed lower connectivity and less tissue-specific expression in Japonica, just as they did in Indica (*≤*0.94× difference, *P≤*6.76×10^-24^; Figure 6D, Supplementary Figure 10D, Supplementary Table 21). Furthermore, these GPTs have lower GC content overall (0.98× difference), although GC content is higher in the 5’ and 3’ UTRs (1.02× difference, *P≤*0.003; Supplementary Figure 10D). The higher UTR GC content contributes to a slightly, but significantly, higher relative N content per transcript (1.003× difference, *P≤*0.001; Supplementary Figure 10D). Furthermore, they are more plastic (1.07× difference), and have lower cross-environment genetic correlations, despite showing higher stability (*≤*0.98× difference, *P≤*0.022; Supplementary Figure 10D). The increased plasticity of GPTs with positive *S* could be contributed to by a denser targeting by small RNAs (1.71× difference, *P*=2.97×10^-6^; Supplementary Figure 10D). Lastly, these transcripts come from loci that show higher gbM (1.18× difference, *P*=0.0007; Supplementary Figure 10D).

Under drought, the influence of many of these factors reversed in direction. GPTs under positive selection for higher expression in dry conditions showed more tissue-specific expression patterns (1.07× difference), and higher connectivity (1.03× difference, *P≤*2.38×10^-6^; Supplementary Figure 10D). They also showed higher overall GC content (1.04× difference, *P*=2.59×10^-11^; Supplementary Figure 10D), which appeared to be driven by higher CDS GC content (1.03× difference), since 5’ UTR GC content was lower (0.97× difference, *P≤*1.82×10^-7^; Supplementary Figure 10D). Furthermore, these transcripts were less plastic than transcripts with negative *S* (0.9× difference, *P≤*0.0003; Supplementary Figure 10D). They were selected for higher rather than lower expression levels (1.03× difference), despite showing lower stability (1.02× difference, *P≤*0.0062; Supplementary Figure 10D). GPTs with positive *S* also showed higher cross-environment environment genetic correlations (1.13× difference), were less densely targeted by small RNAs (0.63× difference), and were transcribed from genes with lower gbM (0.77× difference, *P≤*0.002; Supplementary Figure 10D).

In water-limited conditions selection influenced more features and promoted expression of GPTs with longer CDSs (1.06× difference, *P≤*0.014; Supplementary Figure 10D). Given these observations it is not surprising that these transcripts had higher N requirements as well (1.05× difference), and that the proteins they encode had higher total requirements for N, C and ATP too (*≤* 1.05× difference, *P≤*0.045; Supplementary Figure 10D). Taken together, these observations suggest that, overall, selection may favor higher baseline expression and lower plasticity levels of metabolically costly transcripts and proteins in Japonica. This pattern runs opposite to what we observed for Indica: there, selection favored plasticity of more costly transcripts, and higher baseline expression of less costly transcripts. However, the same factors appear to be important with respect to selection on gene expression for both varietal groups.

### Discrete Plasticity of Translation-Related Transcripts Is Selected Against

No individual transcript made a stringent Bonferroni threshold, but three DPTs involved in essential processes came within an order of magnitude (*P*<0.0001). In Indica, transcripts related to pyrimidine metabolism were enriched among the top transcripts with negative *S* for plasticity, including *OS09T0505800-01* (*S*=-0.42, *P*=5.68×10^-5^; Supplementary Table 22), which codes for a uridine kinase. Pyrimidine metabolism plays key roles in plastid development, allocation of fixed C and abiotic stress responses (Chen and Thelen, 2011; Dong et al., 2019b). In Japonica, selection on transcript level plasticity was strongest, and negative, on *OS02T0633100-02* (*S*=-1.25, *P*=2.41×10^-5^; Supplementary Table 23), coding for the rice ortholog of τ138, which is part of the τB subcomplex of transcription factor IIIC that is involved with recruiting Pol III to transcribe genes encoding tRNAs and other short RNAs (Male et al., 2015). The second transcript under strong negative selection for plasticity was a transcript (*OS10T0430900-01*) from a gene closely related to *OsEDR1*, *OsMAPKKK2* (*S*=-1.16, *P*=2.56×10^-5^; Supplementary Table 23; Rao et al., 2010). In rice, as in other plant species, MAP kinase cascades are at the heart of abiotic and biotic stress responses, as well as growth and developmental programs (Shen et al., 2010).

Local costs of plasticity appeared non-significant, since none of the selection differentials came within an order of magnitude from the Bonferroni threshold (*P*>0.0002; Supplementary Tables 13 and 14), indicating that plasticity costs were most likely to be low at the level of individual transcripts. Combined with the analyses of the distributions of selection differentials, this suggests that costs are more apparent at a polygenic, rather than a monogenic, level.

### Selection Favors Gradual Plasticity of Photosynthesis-Related Transcripts in Japonica

In the Japonica population WGCNA grouped 1,374 GPTs into four transcript co-expression modules, with the remaining 366 transcripts not showing patterns of co-expression (Supplementary Figure 11B, Supplementary Table 26). Out of the four modules, turquoise module 1 showed a signature of significantly elevated levels of positive selection on expression plasticity across environments compared to transcripts in the other modules and the transcripts with unique expression patterns (*S*=0.0059, *P*=0.001), whereas blue module 2 showed a signature of significantly elevated selection for canalization (*S*=-0.0752, *P*<0.0001). Turquoise module 1 was enriched in photosynthesis- and stress response-related GO biological processes (Supplementary Figure 11C), suggesting that selection acted in a polygenic manner on heterogeneity in expression plasticity of photosynthesis- and stress-response related genes across genotypes. Blue module 2 was enriched in translation-related biological processes (Supplementary Figure 11D), which is in keeping with patterns of selection for canalization of translation-related conditionally expressed transcripts (Supplementary Figure 11A, Supplementary Table 24).

Enrichment of TF binding sites in turquoise module 1 for Japonica was very similar to the patterns of enrichment in yellow module 4 for Indica, and transcription factors OsbZIP22, OsE2F1, and OsMYB1R1 again made an appearance (Supplementary Figure 11C). Blue module 2, on the other hand, showed stronger enrichment of TF binding sites for ERF transcription factors, including binding sites for OsSHN1/OsWR1 (Supplementary Figure 11D). This transcription factor in particular has been shown to regulate ribosome components involved in translation at the molecular level (Wilkins et al., 2016), and drought tolerance through altered wax synthesis at the leaf-tissue and whole-plant level (Wang et al., 2012). These overlaps again emphasize the regulatory coherence of turquoise module 1 and blue module 2 and the biological relevance of selection on the plasticity of these modules.

At the level of individual transcripts, two came within an order of magnitude from a Bonferroni cut-off for *P* values on their selection differentials (Supplementary Table 26). One transcript was under selection for expression canalization (*S*=-0.537, *P*=0.0006), and was expressed within blue module 2. This transcript, *OS12T0291400-01*, was generally repressed under drought (log2 fold change of -0.379, FDR *q*6.62×10^-7^), and originated from the gene *OsRBCS5*, encoding a RuBisCO small subunit. Functional OsRBCS5 is necessary for the accumulation of the RuBisCO holoenzyme (Ogawa et al., 2012), and preventing its down-regulation could enhance the robustness of photosynthesis under stress. The other transcript was not co-expressed within a module and was under positive selection for expression plasticity (*S*=0.334, *P*=0.0006). This was *OS06T0602700-00*, coding for the plastidic adenine nucleotide uniporter OsBT1-3 (Hu et al., 2017). Expression of *OsBT1-3* appears to be affected by drought in some, but not all, accessions (environment FDR *q*=0.318), and is tied to chloroplast development and the number of chloroplasts in leaves (Hu et al., 2017).

## Supplementary Figures

**Supplementary Fig. 1.**
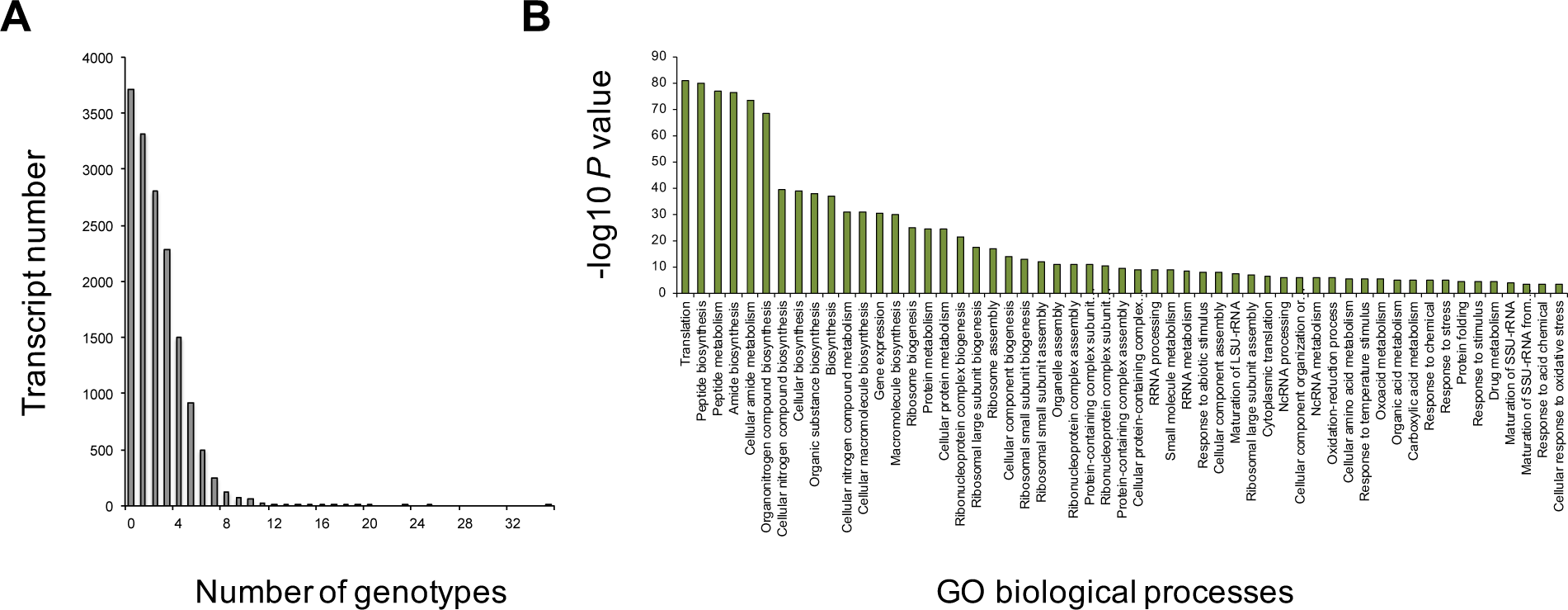
Population-level heterogeneity in transcriptional plasticity. **a,** Number of Japonica genotypes in which each transcript is influenced significantly by drought (*P*<0.05) for each transcript identified from one-way ANOVAs. **b,** Gene ontology biological processes enriched in the tail of the distribution of differentially expressed transcripts with each transcript influenced by drought in at least 10 genotypes (FDR *q*<0.05).

**Supplementary Fig. 2.**
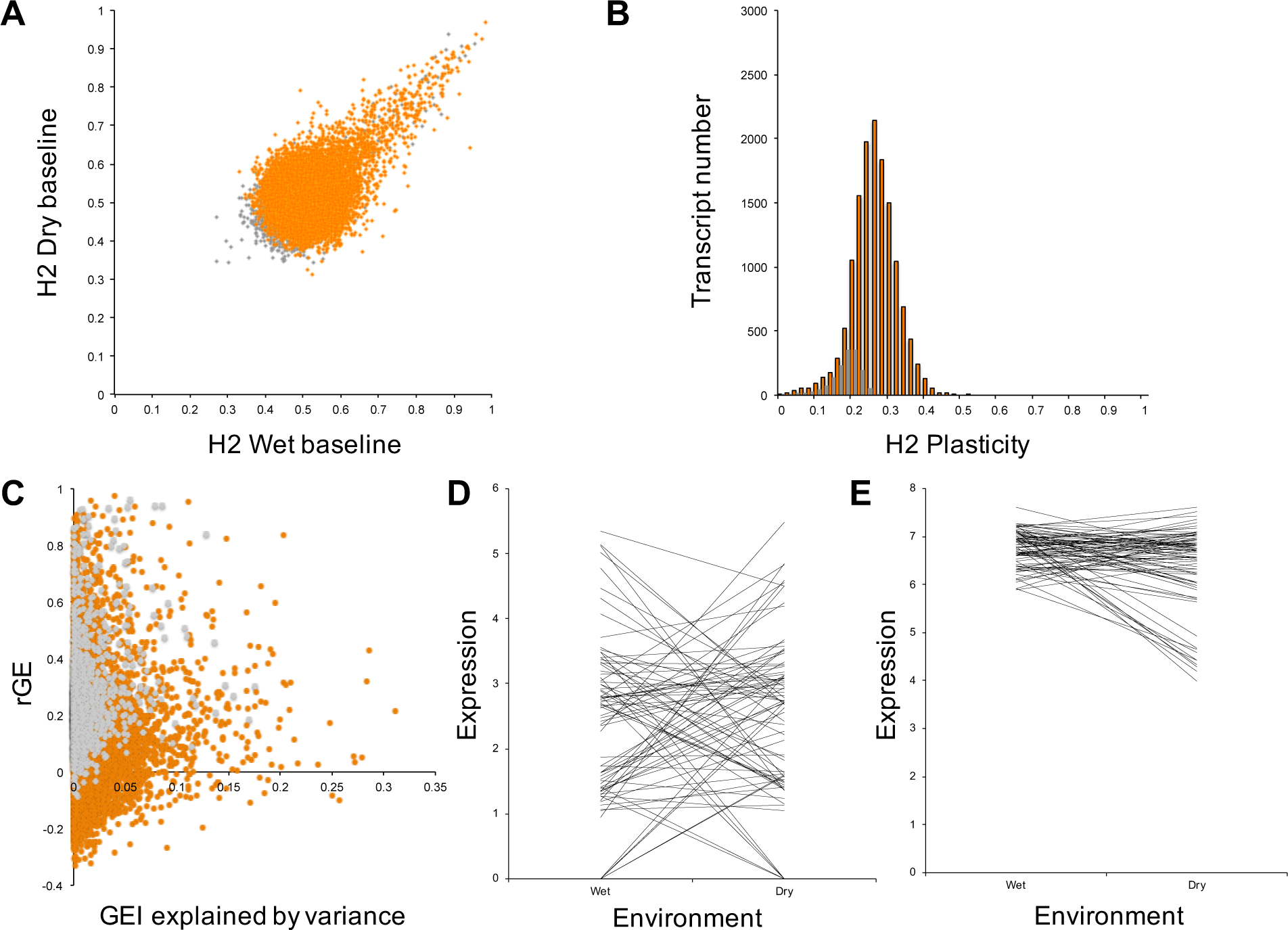
Heritability of transcript level plasticity. **a,** Bivariate plot of broad-sense heritability (*H^2^*) estimates for baseline transcript levels in the Japonica populations in wet and dry conditions. Orange dots indicate significant G×E interaction variance (FDR *q*<0.001), and grey non-significant. **b,** Distribution of *H^2^* estimates for transcript level plasticity across wet and dry conditions. Orange columns denote significant G×E interaction variance (FDR *q*<0.001), and grey non-significant. **c,** Proportion of G×E interaction variance attributed to changes in among-accession variance in wet and dry conditions versus cross-environment genetic correlation. Orange dots indicate significant G×E interaction variance (FDR *q*<0.001), and grey non-significant. **d,** Reaction norms of a transcript for which G×E interaction variance is mostly determined by deviation of cross-environment genetic correlation from unity, as indicated by abundant line crossing. **e,** Reaction norms of a transcript for which more G×E interaction variance is determined by changes in among-accession variance in wet and dry conditions, as indicated by less-abundant line crossing and wider among-accession variance in one environment than the other.

**Supplementary Fig. 3.**
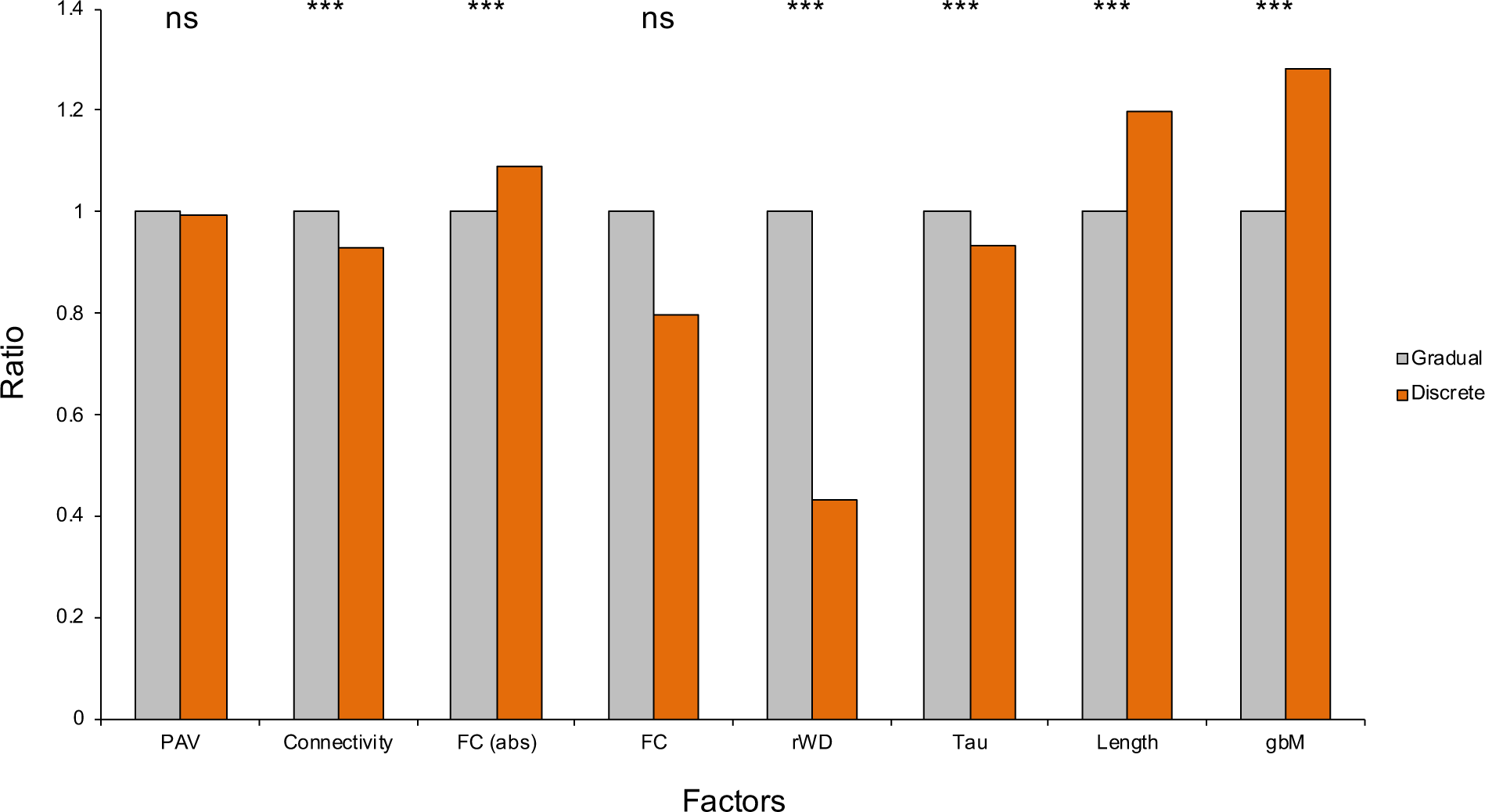
Structural and regulatory features of gradually and discretely plastic transcripts. **a,** Average levels of factors that characterize discretely plastic transcripts (DPTs) relative to average levels of these factors for gradually plastic transcripts (GPTs) in Japonica. Values for the former have been normalized relative to the latter. PAV = presence/absence variation, FC = fold change, Tau = tissue specificity, gbM = gene body methylation, ns = non-significant; *** indicates *P*<0.001.

**Supplementary Fig. 4.**
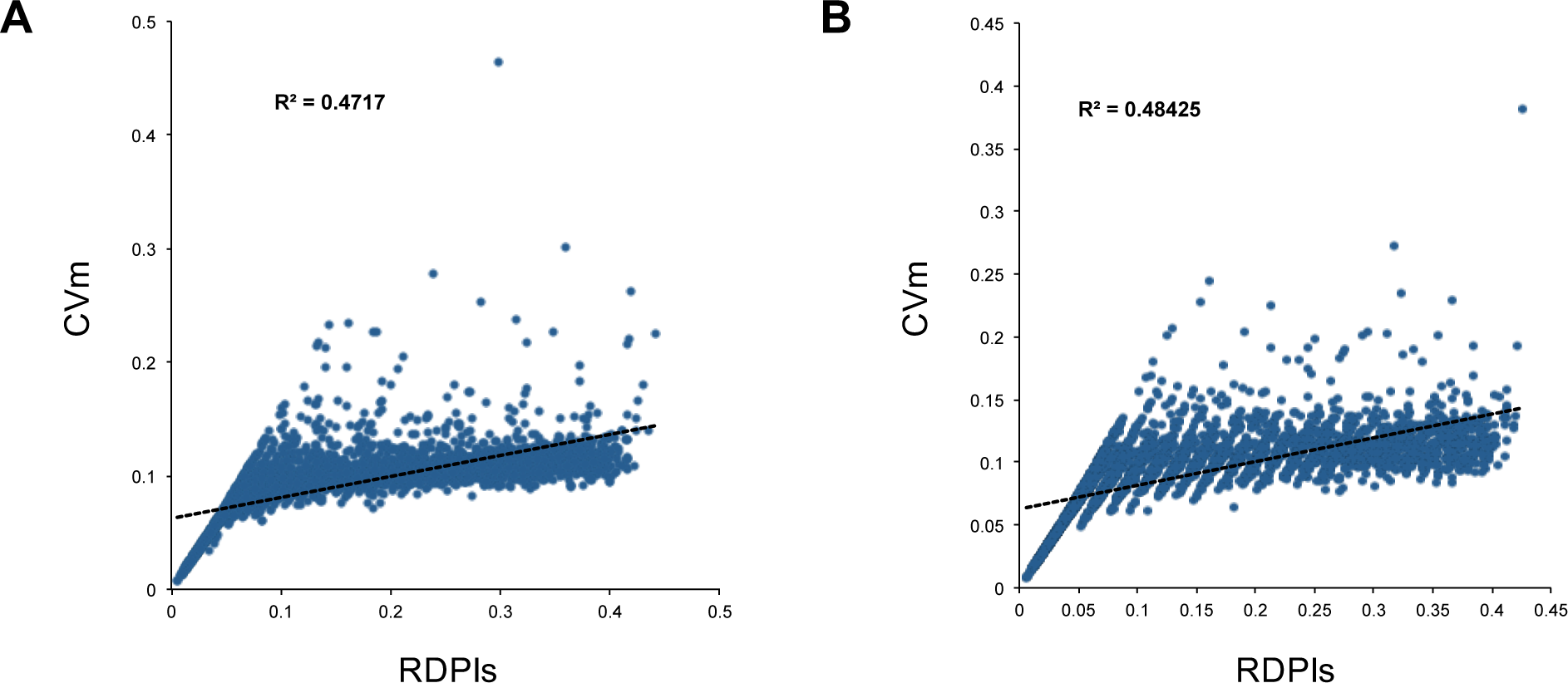
Correlation between different plasticity indices for gradually plastic transcripts. **a,** In the Indica populations values of the simplified relative distance plasticity index (RDPI_s_), which was used to estimate plasticity of gradually plastic transcripts (GPTs), show significant correlation with values of the coefficient of variation over the environments based on means (CV_m_) calculated on the same transcripts. This ensures that the latter index can be used for estimating plasticity of discretely plastic transcripts (DPTs). **b,** The same held true for the Japonica populations.

**Supplementary Fig. 5.**
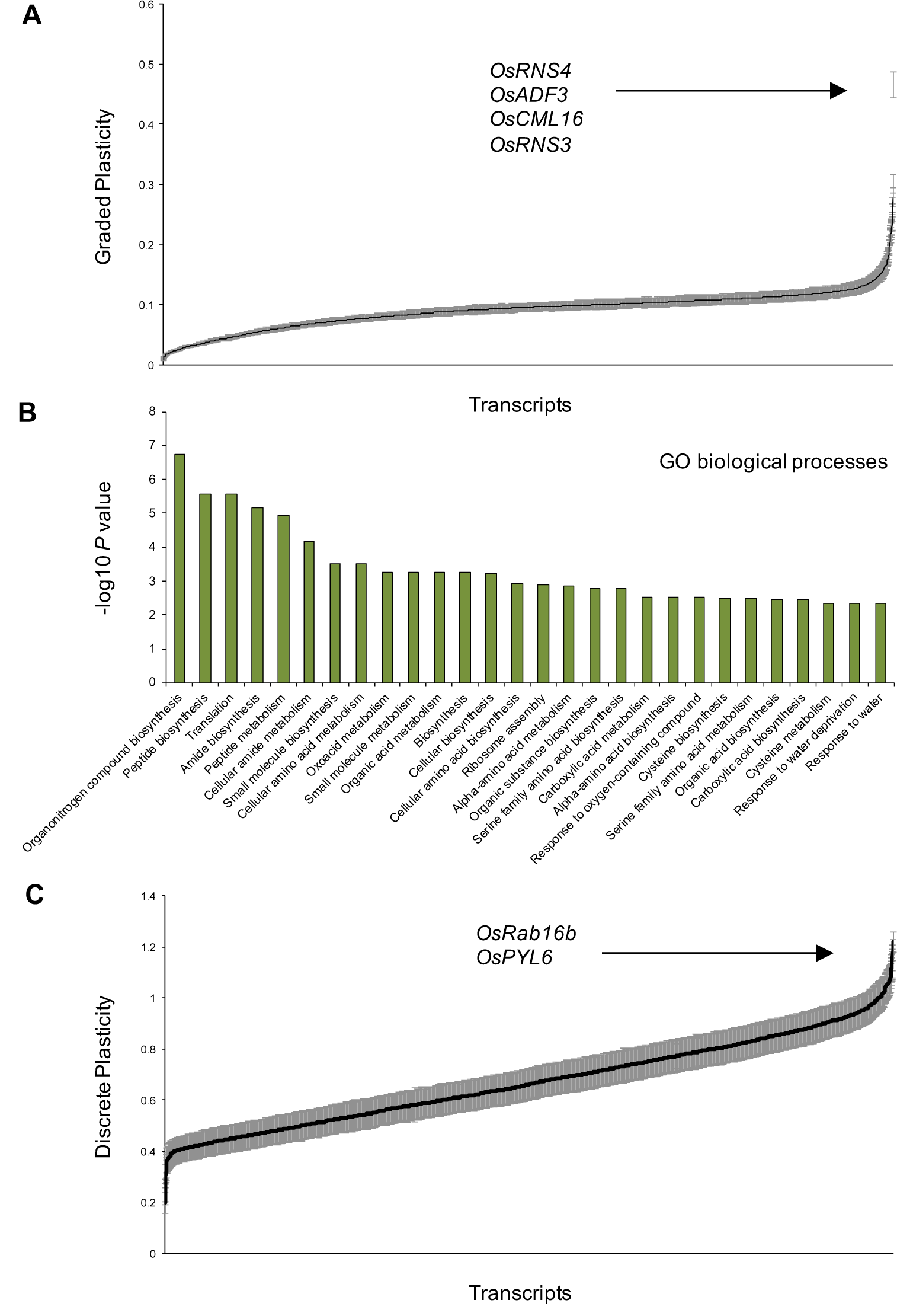
Biological characterization of gradually and discretely plastic transcripts in Indica. **a,** Distribution of plasticity levels for gradually plastic transcripts (GPTs). **b,** Gene ontology biological processes enriched in the high-value tail of the plasticity distribution for GPTs (FDR *q*<0.05). **c,** Previously characterized drought stress-related transcripts in the high-value tail of the plasticity distribution for discretely plastic transcripts (DPTs).

**Supplementary Fig. 6.**
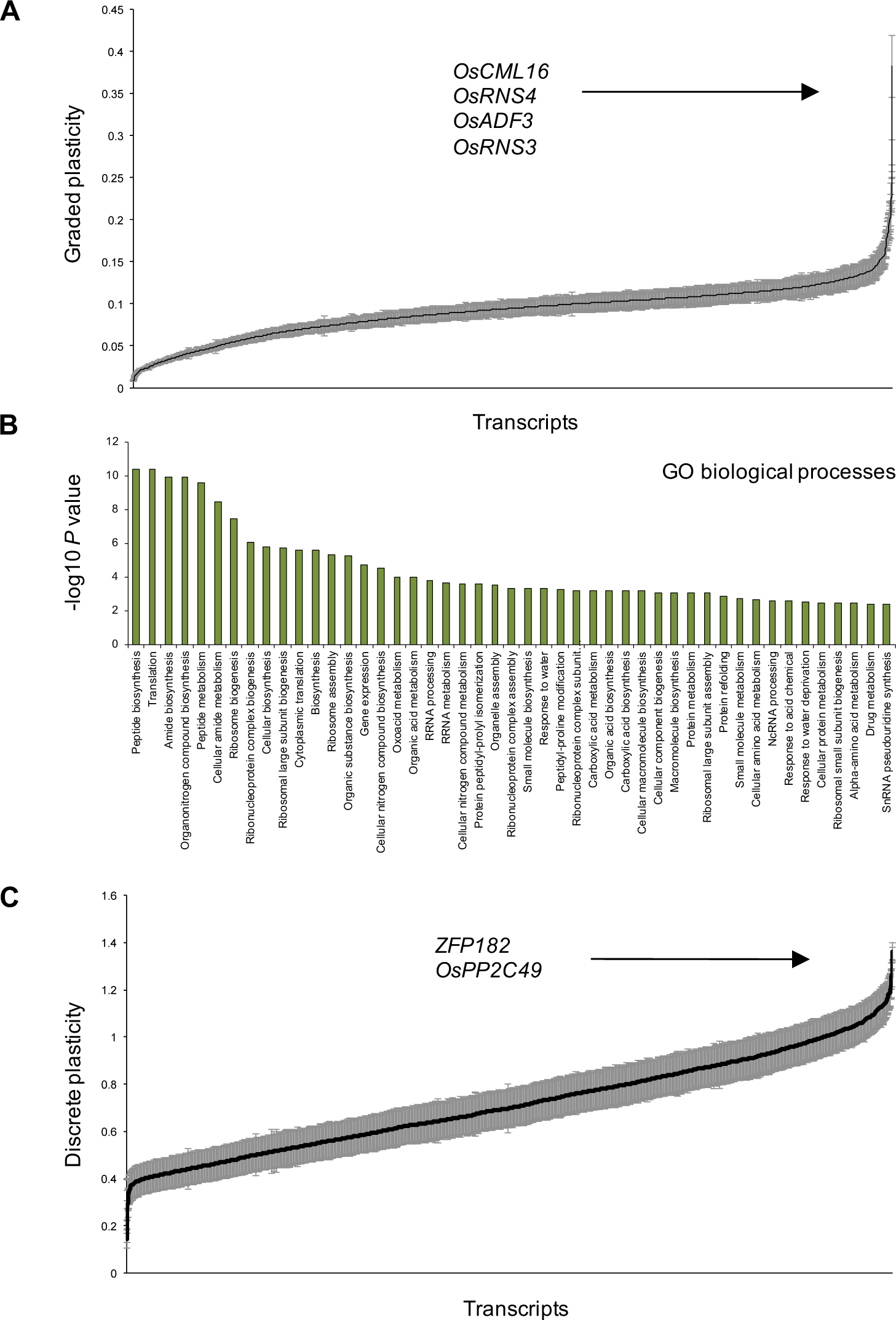
Biological characterization of gradually and discretely plastic transcripts in Japonica. **a,** Distribution of plasticity levels for gradually plastic transcripts (GPTs). **b,** Gene ontology biological processes enriched in the high-value tail of the plasticity distribution for GPTs (FDR *q*<0.05). **c,** Previously characterized drought stress-related transcripts in the high-value tail of the plasticity distribution for discretely plastic transcripts (DPTs).

**Supplementary Fig. 7.**
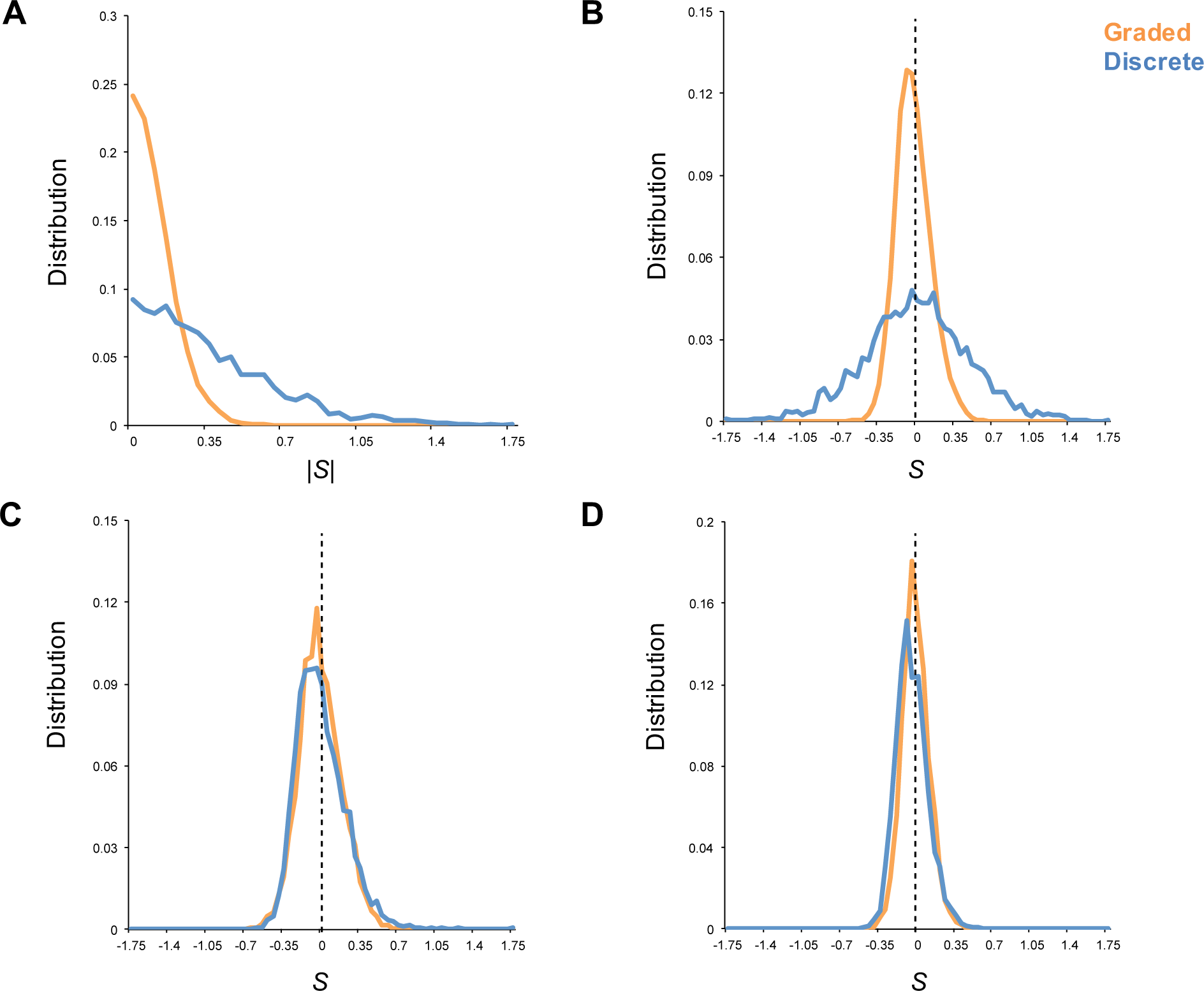
The strength and pattern of selection on expression plasticity differs between gradually and discretely plastic transcripts. **a,** The strength of selection |*S*| on gene expression plasticity in Japonica when considering across-environment (global) fecundity as fitness measure differed between gradually plastic (orange) and discretely plastic (blue) transcripts (GPTs and DPTs, respectively). **b,** Positive directional selection was in general similarly strong as negative directional selection for the plasticity of both GPTs and DPTs. **c,** Positive directional selection on plasticity generally outweighed plasticity costs for GPTs in wet conditions. **d,** This pattern was similar under drought. For DPTs selection was biased towards plasticity costs in dry conditions, and this pattern appeared the exact opposite in wet conditions. See supplementary text for statistical information.

**Supplementary Fig. 8.**
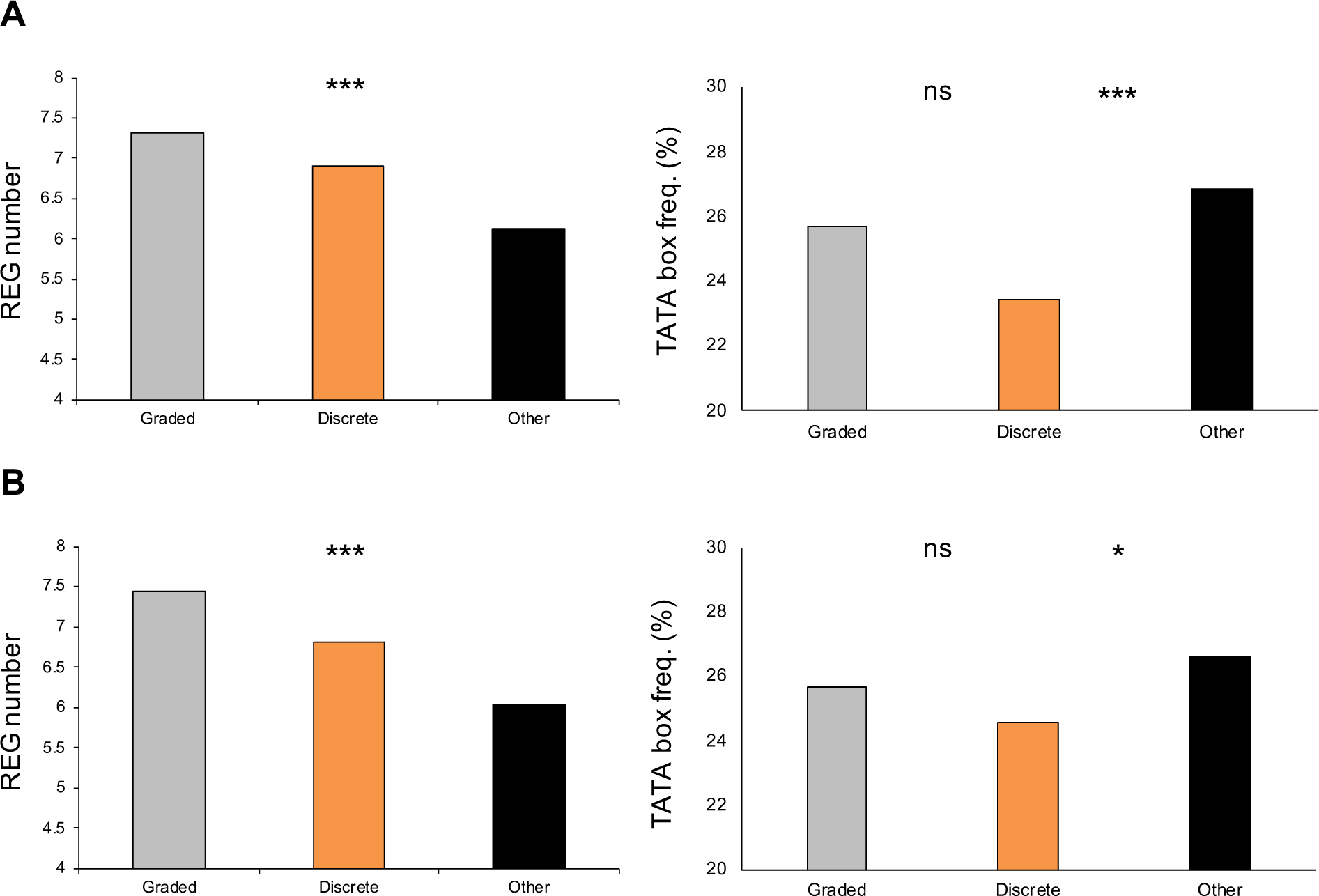
Characterization of gradually and discretely plastic transcripts. **a,** Gradually and discretely plastic transcripts (GPTs and DPTs, respectively) in Indica differed significantly in the frequency with which TATA boxes occur in their genes’ promoters with respect to the frequency of TATA boxes in the promoters of other leaf-expressed genes (Fisher’s exact test; one-tailed *P*=0.127 for GPTs vs. other transcripts, and one-tailed *P*=0.001 for DPTs vs. other transcripts, respectively), and differed in the numbers of *cis*-regulatory promoter elements (REGs) in their genes’ promoters as well (one-way ANOVA; *P*<0.0001, lower 95% CI for GPTs higher than transcriptome-wide average REG number of 6.65 [6.98, 7.66], which was within 95% CI for DPTs [6.5, 7.31]). **b,** We observed similar patterns across the Japonica populations for the frequency of TATA boxes (Fisher’s exact test; one-tailed *P*=0.173 for GPTs vs. other transcripts, and one-tailed *P*=0.027 for DPTs vs. other transcripts, respectively), and REG numbers in promoters (one-way ANOVA; *P*<0.0001, lower 95% CI for GPTs higher than transcriptome-wide average REG number of 6.65 [7.1, 7.77], which was within 95% CI for DPTs [6.43, 7.18]). Ns = non-significant; *** indicates *P*<0.001; * indicates *P*<0.05.

**Supplementary Fig. 9.**
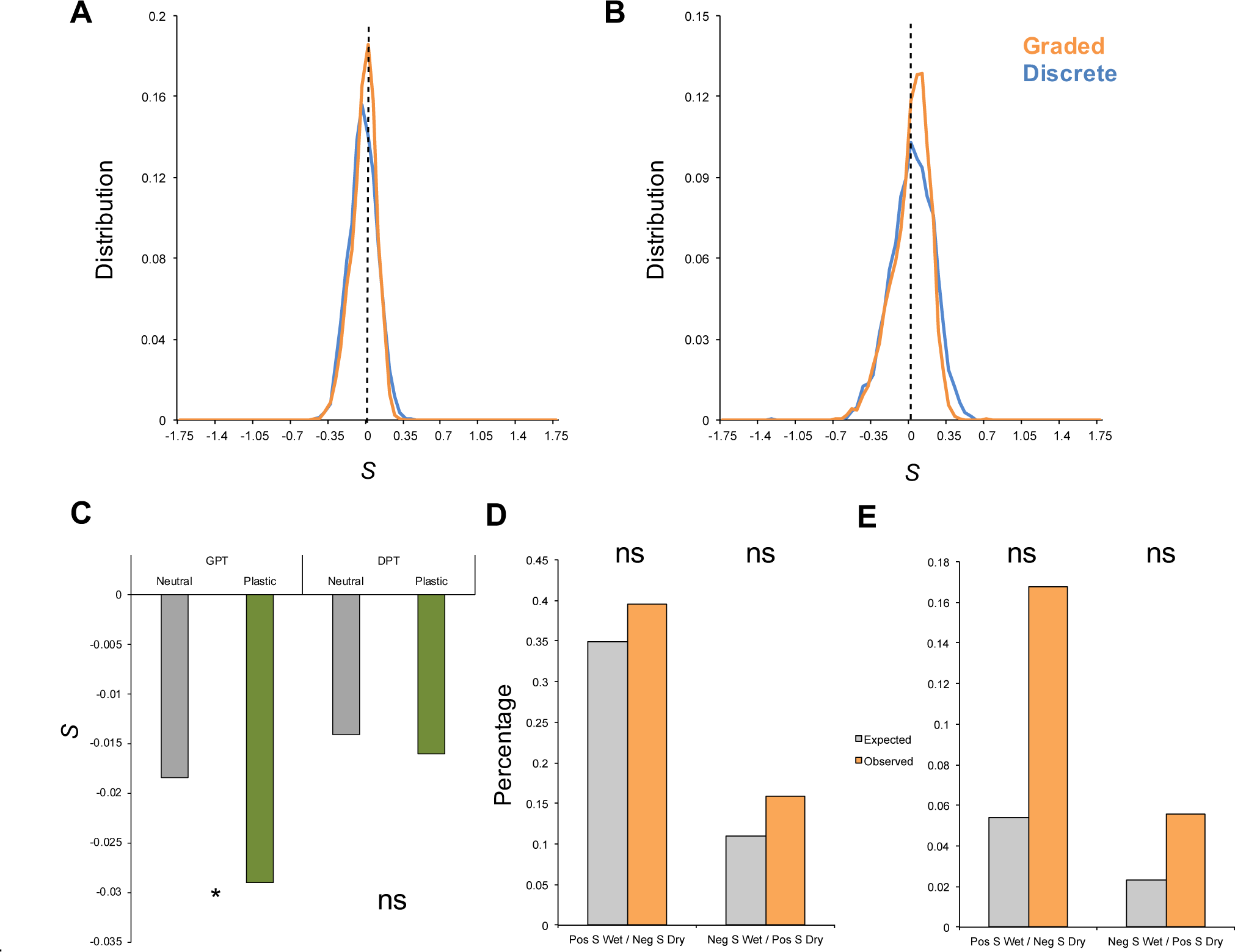
The strength and pattern of selection on baseline expression levels tend to be congruent between gradually and discretely plastic transcripts. **a,** Under drought selection tended to benefit higher baseline expression levels for both gradually and discretely expressed transcripts (GPTs and DPTs, respectively) in Japonica. **b,** This pattern held true in wet conditions as well. See supplementary text for statistical information. **c,** Selection on baseline expression under drought was countergradient with the levels of expression plasticity. **d,** Antagonistic pleiotropy is as common as expected by chance for GPTs. **e**, For DPTs antagonistic pleiotropy was as common as expected by chance as well. Ns = non-significant; * indicates *P*<0.05.

**Supplementary Fig. 10.**
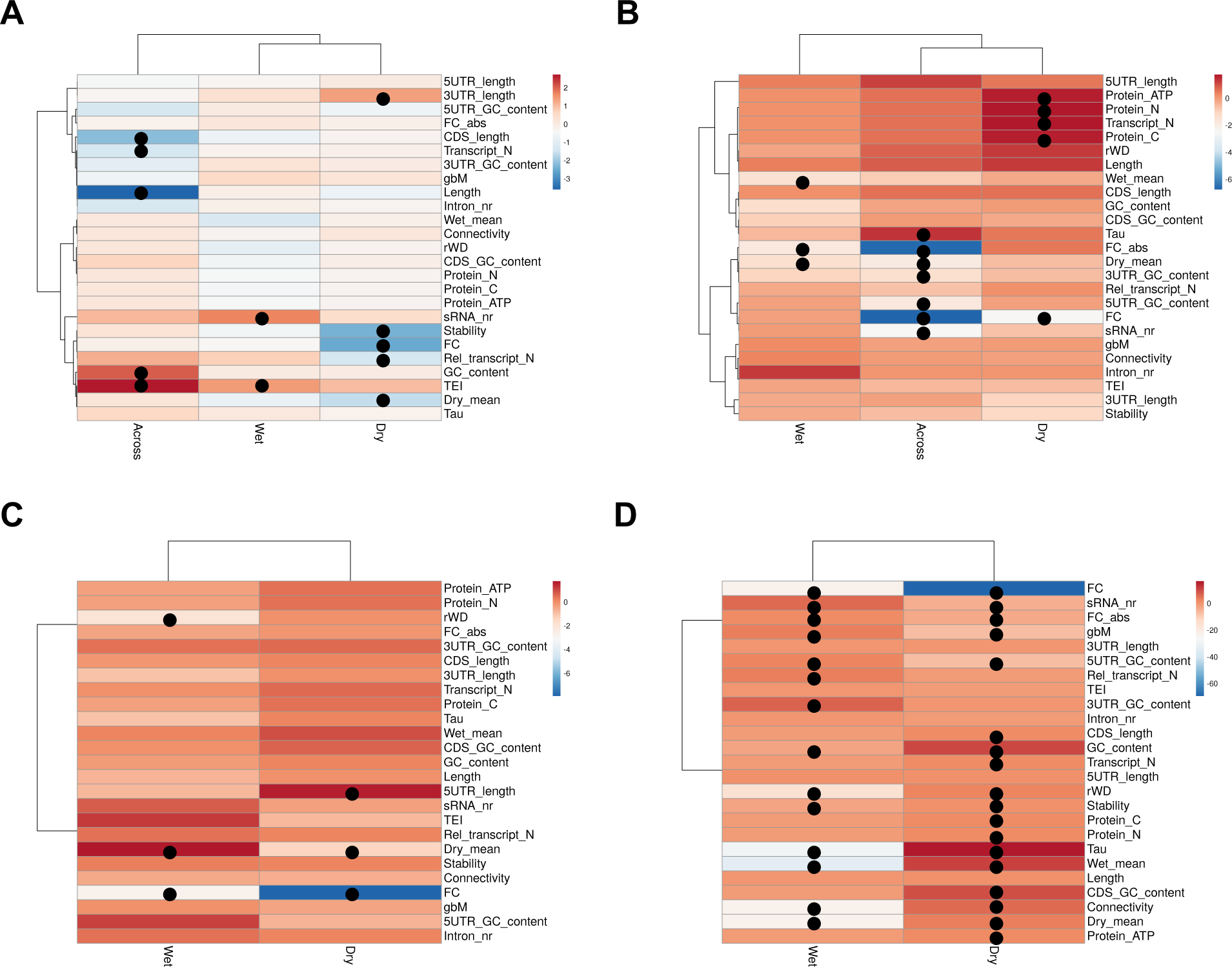
Comparison of defining characteristics of transcripts with positive and negative selection differentials on expression plasticity and baseline expression in the Japonica populations. Gradually and discretely plastic transcripts (GPTs and DPTs, respectively) were compared for 25 architectural, biochemical and network characteristics in the five selection scenarios we studied for DPTs and GPTs separately: across- and within-environment selection on plasticity for DPTs **(a)** and GPTs **(b)**, as well as selection on baseline expression in wet and dry conditions for DPTs **(c)** and GPTs **(d)**. Panels depict the negative log10 of the *P* values of the *t*-tests between the transcripts with positive and negative *S*. Positive changes indicate a characteristic was more strongly associated with transcripts defined by positive *S* than with transcripts defined by negative *S*, while negative changes indicate the opposite pattern.

**Supplementary Fig. 11.**
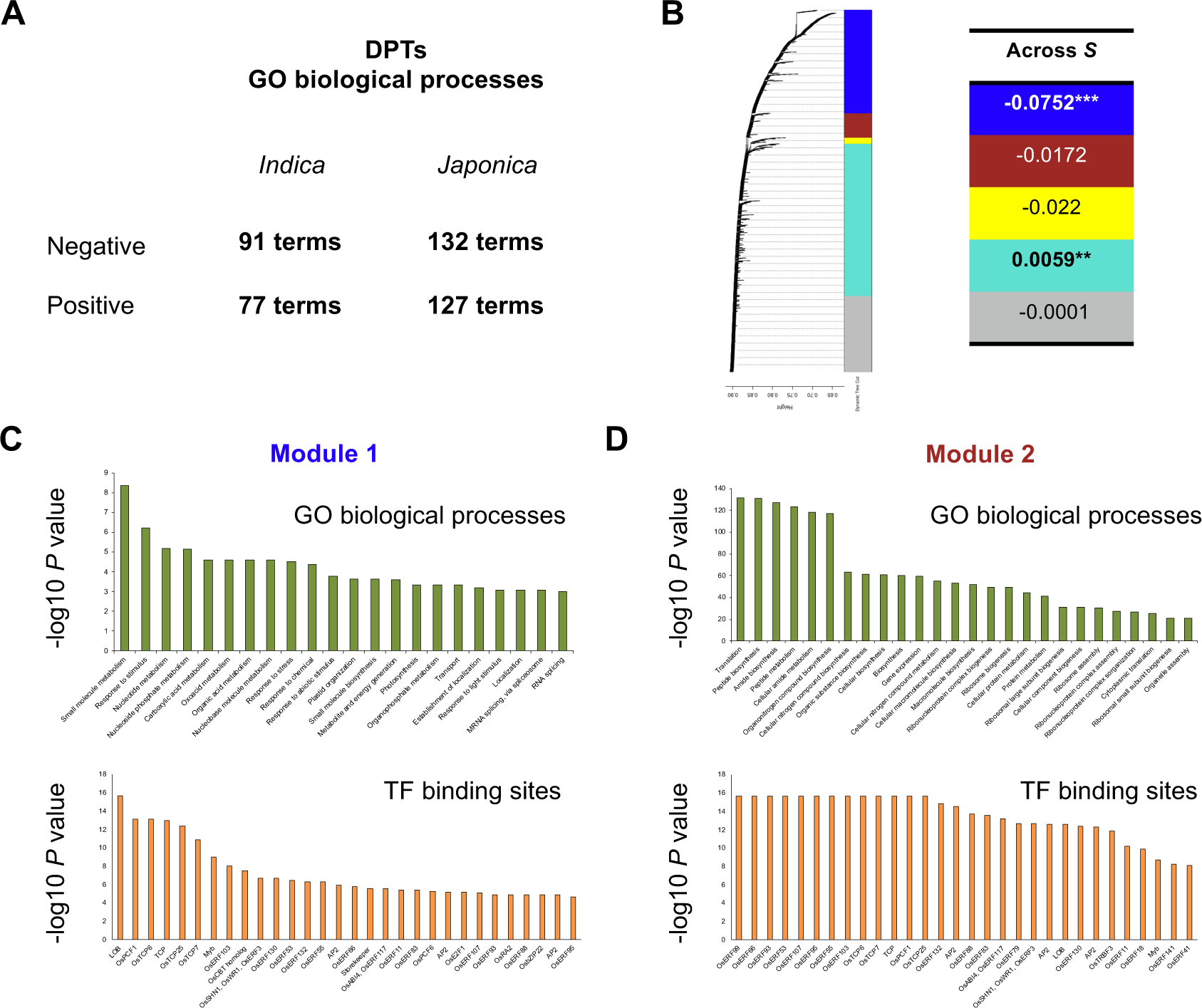
Selection may have polygenic effects on plasticity levels of transcript co-expression modules. **a,** Discretely plastic transcripts in Japonica are enriched for different Gene Ontology (GO) biological processes among transcripts with positive and negative selection differentials on their plasticity levels. **b,** Selection acts more strongly on two modules of co-expressed gradually plastic transcripts than on other modules. See supplementary text for statistical information. **c,** The modules under selection are enriched for photosynthesis-related and translation-related biological processes, respectively. **d,** In addition, the gene promoters of the first module’s transcripts show enrichment of binding sites for bZIP, MYB, and TCP transcription factors, whereas the gene promoters of the second module’s transcripts show enrichment of binding sites for AP2 ERF transcription factors.

## Supplementary Tables

Supplementary Table 1 | One-way analysis of variance in the transcriptome of each Indica accession across wet and dry field conditions.

Supplementary Table 2 | Gene set enrichment analysis of transcripts showing environmentally biased expression patterns in Indica accessions.

Supplementary Table 3 | One-way analysis of variance in the transcriptome of each Japonica accession across wet and dry field conditions.

Supplementary Table 4 | Gene set enrichment analysis of transcripts showing environmentally biased expression patterns in Japonica accessions.

Supplementary Table 5 | Analysis of genotype-environment interaction in the transcriptome of the Indica populations in wet and dry field conditions.

Supplementary Table 6 | Analysis of genotype-environment interaction in the transcriptome of the Japonica populations in wet and dry field conditions.

Supplementary Table 7 | Metadata of factors that may characterize transcripts with graded and discrete plasticity in the Indica populations

Supplementary Table 8 | Metadata of factors that may characterize transcripts with graded and discrete plasticity in the Japonica populations

Supplementary Table 9 | Gene set enrichment analysis of transcripts showing graded and discrete plasticity in the Indica populations.

Supplementary Table 10 | Gene set enrichment analysis of transcripts showing graded and discrete plasticity in the Japonica populations.

Supplementary Table 11 | Between- and within-environment selection differentials for plasticity of gradually plastic transcripts in the Indica populations.

Supplementary Table 12 | Between- and within-environment selection differentials for plasticity of gradually plastic transcripts in the Japonica populations.

Supplementary Table 13 | Between- and within-environment selection differentials for plasticity of discretely plastic transcripts in the Indica populations.

Supplementary Table 14 | Between- and within-environment selection differentials for plasticity of discretely plastic transcripts in the Japonica populations.

Supplementary Table 15 | Metadata of gene regulatory network factors that may characterize transcripts with graded and discrete plasticity

Supplementary Table 16 | Local selection differentials for baseline expression of gradually and discretely plastic transcripts in the Indica populations.

Supplementary Table 17 | Local selection differentials for baseline expression of gradually and discretely plastic transcripts in the Japonica populations.

Supplementary Table 18 | Cogradient selection and Conditional Neutrality / Antagonistic Pleiotropy (CNAP) analyses for the Indica populations.

Supplementary Table 19 | Cogradient selection and Conditional Neutrality / Antagonistic Pleiotropy (CNAP) analyses for the Japonica populations.

Supplementary Table 20 | Gene architectural, metabolic and regulatory network factors that are under differential selection the Indica populations.

Supplementary Table 21 | Gene architectural, metabolic and regulatory network factors that are under differential selection the Japonica populations.

Supplementary Table 22 | Gene set enrichment analysis of discretely plastic transcripts under selection in the Indica populations.

Supplementary Table 23 | Gene set enrichment analysis of discretely plastic transcripts under selection in the Japonica populations.

Supplementary Table 24 | Overlap in enriched gene sets of discretely plastic transcripts under selection in the Indica and Japonica populations.

Supplementary Table 25 | Co-expression modules of gradually plastic transcripts in the Indica populations.

Supplementary Table 26 | Co-expression modules of gradually plastic transcripts in the Japonica populations.

